# The gene expression signature of electrical stimulation in the human brain

**DOI:** 10.1101/2023.09.21.558812

**Authors:** Snehajyoti Chatterjee, Muhammad Elsadany, Yann Vanrobaeys, Annie I Gleason, Brian J. Park, Shane A. Heiney, Ariane E. Rhone, Kirill V. Nourski, Budhaditya Basu, Utsav Mukherjee, Lucy Langmack, Christopher K. Kovach, Zsuzsanna Kocsis, Yukiko Kikuchi, Yaneri A. Ayala, Mark Bowren, Marco M. Hefti, Ethan Bahl, Kailin Yang, Jeremy D. Greenlee, Hiroto Kawasaki, Hiroyuki Oya, Matthew A. Howard, Christopher I. Petkov, Thomas Nickl-Jockschat, Li-Chun Lin, Jacob J Michaelson, Ted Abel

## Abstract

Direct electrical stimulation (eSTIM) is widely used clinically, from neurosurgical mapping to therapeutic interventions for neurological and neuropsychiatric disorders^1–10^. Despite over a century of application, its molecular and cellular underpinnings remain unknown. Here, using state-of-the-art single-nuclei multiomic profiling, we map changes in cell-type-specific gene expression and chromatin accessibility *in vivo* in the human cortex following eSTIM of neurosurgery patients. eSTIM impacts a network of cells that extends beyond excitatory neurons to include inhibitory neurons, astrocytes, oligodendrocytes and microglia. We observed an upregulation of canonical immediate-early genes (IEGs: *FOS*, *NPAS4*, *EGR4*) in excitatory and inhibitory neurons and induction of cytokine-related genes *CCL3 and CCL4* in microglia. The cross-species conservation of this gene signature, together with our examination of a cohort of both epilepsy and cancer patients, underscores the fundamental role of these changes in stimulation-driven plasticity while controlling for disease and environmental confounds. Our study of changes in chromatin accessibility reveals a common code that involves a cell-type specific signature of transcription factor binding motifs for members of the EGR family. By addressing these previously unexplored questions about activity-induced gene expression *in vivo* in the human brain, our findings challenge the long-standing neuron-centric view of eSTIM, highlighting the broader role of non-neuronal cells, including microglia, in mediating the impact of brain stimulation.

## Main

Electrical stimulation of the human brain has become an indispensable clinical tool for diagnosis and therapy^1–8^. In neurosurgical patients, cortical stimulation is the gold standard diagnostic tool to identify the location of functionally relevant brain regions that are critical for speech, language, or motor function^11,12^ to avoid potential damage to these regions during surgery. eSTIM protocol that is commonly employed to either induce movement when motor cortex^9^ is stimulated or lead to speech arrest^10^ when language eloquent cortex is stimulated. Transient changes in gene expression may support the ability of the brain to adapt to changes in the environment or to respond to stimuli such as deep brain stimulation (DBS)^13^. Dynamic transcriptomic patterns are essential for cognition^14–16^, affective processing^17^, addiction^18^, and the initiation of behaviors^19,20^, and eSTIM may in part exert effects via alterations in gene expression. Although diagnostic and therapeutic brain stimulation is conducted daily in thousands of patients worldwide, the molecular impact of eSTIM in the human brain remains unknown. Recent advances in molecular sequencing techniques have revolutionized transcriptomics by enabling mapping of changes in transcription and chromatin accessibility in single cells^21^. However, these experiments have only been conducted in rodents^22,23^ and neurons derived from human induced pluripotent stem cells^24^. A recent pioneering study utilized a single nuclei molecular approach to link human brain transcriptomics signatures with oscillatory signatures of memory consolidation in epileptic patients undergoing an episodic memory task, but these gene expression measures were obtained many days after assessment of intracranial recording data during the task^25,26^. Therefore, research is needed to evaluate the immediate effects of eSTIM on single-cell gene expression in the living human brain. Studies investigating gene expression profiles after eSTIM are particularly difficult to perform in human brains due to the requirement that tissue is sampled at precise temporal windows before and after eSTIM.

Here, in epilepsy and cancer patients undergoing resection in the temporal or frontal lobe, samples were obtained before and after experimental electrical stimulation. By analyzing gene expression from bulk tissue at baseline and minutes after eSTIM from the same patient, along with our analysis of samples taken from patients that did not receive eSTIM, we are able to distinguish genes responsive to eSTIM from the genes altered nonspecifically by surgical or disease-based factors that would be stable across samples. Our parameters for eSTIM were based on those regularly used in clinical mapping^27,28^; moreover, we referenced the human results to a similar eSTIM paradigm in mice, revealing a similar transcriptional profile. Finally, using a single-nuclei multi-omics approach, we provide insight into the cell-type-specific transcriptomic and epigenomic responses to eSTIM in the human brain. eSTIM leads to robust, transcriptomic changes in activity-dependent canonical immediate-early genes (IEGs) in the excitatory and inhibitory neuronal clusters, while, microglia exhibited the greatest changes, including robust induction of cytokine-related genes. This study provides fundamental insights into changes in cell-type specific molecular signatures in the human cortex after eSTIM, laying the groundwork for a molecular understanding of the impact of this fundamental tool in clinical neurosurgery, diagnosis, and treatment.

## Results

Initially, we recruited eight adult neurosurgical patients undergoing surgical resection of seizure foci following clinical monitoring to treat epilepsy. The patients provided informed consent to participate in this research and were informed that tissue samples would only be taken from tissue that would require resection for clinical treatment. The epileptic participants underwent an anterior temporal lobectomy (ATL) for access to a medial temporal lobe epileptogenic site, during which samples were resected from the neocortex and processed immediately after removal. The participants were distributed between two experimental paradigms. In the first group with the electrically stimulated paradigm (4 participants), a baseline tissue sample was resected from the ATL after the craniotomy and durotomy, exposing the temporal lobe. Then, an adjacent region of the cortex was stimulated using a stimulation protocol commonly used for bipolar electrical stimulation (50 Hz) for 2 minutes^27,28^, and resected approximately thirty minutes later (27.86 ± 10.93 min), a sample was taken from the stimulated region **(Fig. 1a; Supplementary Fig. 1a-c, Supplementary Table 1).** In the second group with the unstimulated paradigm (4 participants), a sample was taken at baseline, and then a second sample was taken without stimulation about 30 minutes later (37.3 ± 12.5 mins, **Fig. 1b; Supplementary Fig. 1a-c, Supplementary Table 1)**. The pre-stimulation sample and the unstimulated paradigm groups serve as controls for changes in baseline gene expression and changes in gene expression occurring as a result of the surgical procedure. All the resected tissue samples from this study, from both the stimulated and unstimulated paradigms were taken well outside the seizure focus (27.3 ± 5.6 mm from the border of the seizure focus**, Supplementary Fig. 1b**).

**Figure 1.**
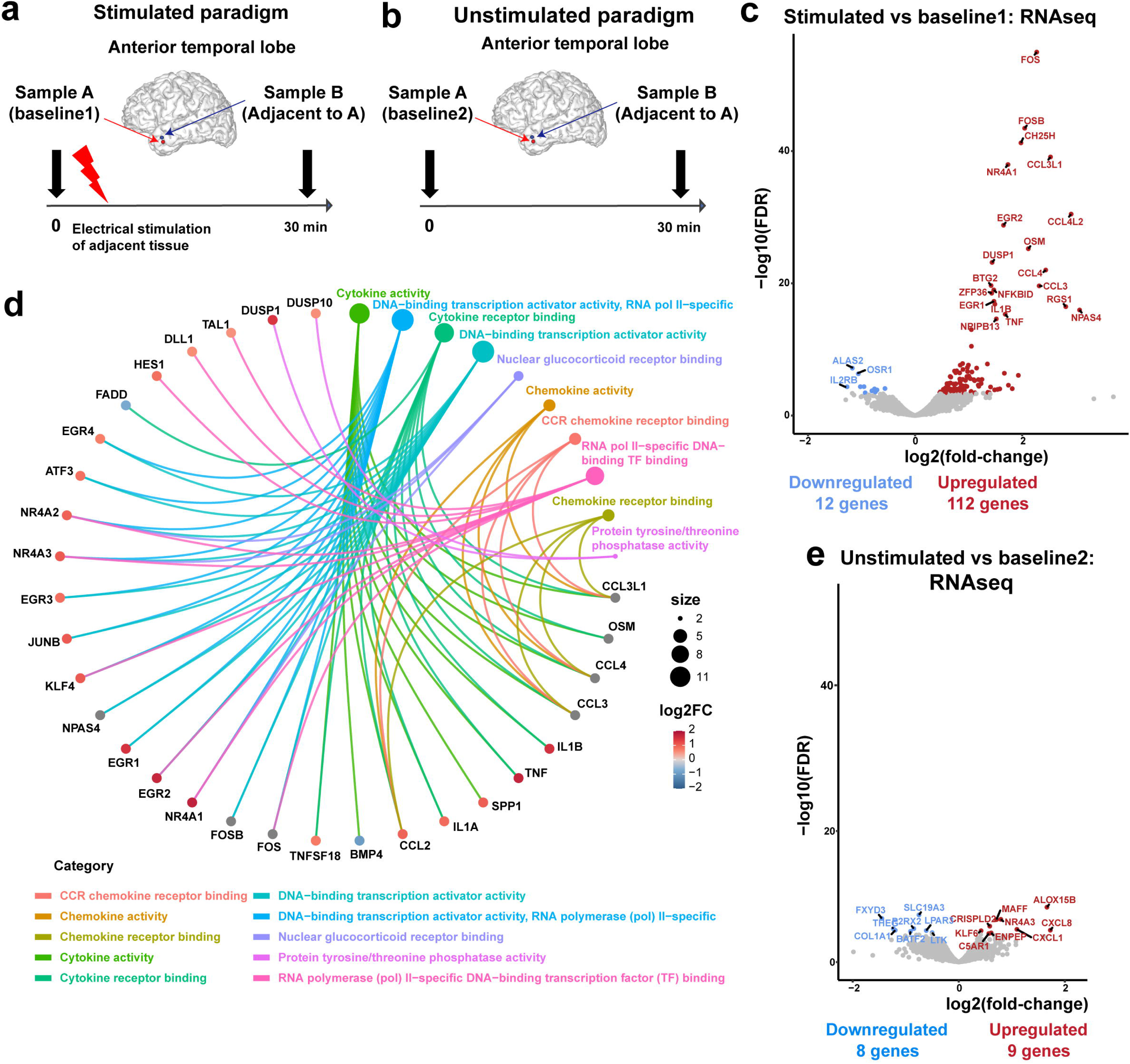
Electrical stimulation (eSTIM) of the human cortex induces changes in genes associated with cytokine activity and transcription regulation. **a, b.** Schematics of stimulated (**a**) and unstimulated (**b**) paradigms in the human anterior temporal lobe. For both paradigms sample A was taken at T = 0 minutes then an adjacent sample B was taken at either 30 min after stimulation (**a**, stimulated paradigm, 4 participants) or 30 min after sample A was taken (**b**, unstimulated paradigm, 4 participants). **c**. Volcano plot showing gene expression changes in the stimulated samples. The most significant genes (FDR < 0.05) are labeled in red (upregulated) or blue (downregulated). **d**. Cnet plot showing pathway enrichment analysis of the genes significantly (FDR < 0.05) differentially expressed in the stimulated samples. **e**. Volcano plots showing gene expression changes in the unstimulated samples. The most significant genes (FDR < 0.05) are labeled in red (upregulated) or blue (downregulated).

Tissue samples were subjected to bulk whole-transcriptome RNA sequencing (RNA-seq) to identify differentially expressed genes following eSTIM. Bulk RNA-seq analysis from the stimulated and corresponding baseline samples revealed 124 differentially expressed genes following eSTIM, with 112 up-regulated and 12 down-regulated genes **(Fig. 1c**; **Supplementary Table 2)**. The deconvolution of the bulk RNA-seq data additionally revealed comparable proportions of cell types between the baseline and stimulated samples (**Supplementary Fig. 2**). Enrichment network analysis was used to identify the pathways most represented among the differentially expressed genes in the stimulated paradigm. The top significant pathways were enriched with genes involved in cytokine activity, DNA-binding transcription activator activity (RNA Pol II), cytokine receptor binding, DNA-binding transcription activator activity, nuclear glucocorticoid receptor binding, chemokine activity, CCR chemokine receptor binding, RNA pol II specific DNA binding, chemokine receptor binding, and protein phosphatase activity **(Fig. 1d).**

Bulk RNA-seq analysis from the unstimulated paradigm comparing unstimulated samples with corresponding baseline samples identified differential expression of only 17 genes, with nine up-regulated and eight down-regulated genes **(Fig. 1e, Supplementary Table 3).** Only one gene, *NR4A3*, was found to be differentially expressed in both the stimulated and unstimulated groups. The lack of overlap between the differentially expressed genes observed in our stimulated and unstimulated groups suggests that the changes in gene expression that we see following electrical stimulation do not reflect disease state or surgical effects such as craniotomy, brain temperature, and anesthesia, but reflect changes due to eSTIM.

We used a similar eSTIM paradigm in the mouse non-primary auditory cortex to investigate whether these gene expression changes are unique to human samples and to determine whether they are related to disease state ^29,30^. Bulk RNA-seq was performed on samples collected 30 minutes following electrical stimulation and unstimulated samples collected at the same time from the contralateral side (**Fig. 2a**). Bulk RNA-seq identified 44 upregulated and 107 downregulated genes (**Fig. 2b, Supplementary Table 4**). Pathway analysis identified from the upregulated genes in the mouse was enriched for cytokine activity, chemokine activity, cytokine receptor binding, and CCR chemokine receptor binding (**Fig. 2c**). We also found upregulated pathways linked to transcription and post-transcription regulatory pathways, such as mRNA 3’-UTR binding, mRNA3’-UTR AU-rich region binding, and transcription co-repressor activity. To assess the conservation of transcriptional responses, we performed a Rank-Rank Hypergeometric Overlap (RRHO) analysis^31,32^ comparing the bulk RNA-seq differential expression results from stimulated human samples to those from stimulated mouse samples. This analysis identified a statistically significant concordance in upregulated genes between the two species. The RRHO test revealed 32 genes that were consistently upregulated in both human and mouse following electrical stimulation, highlighting a partially conserved transcriptional program (**Fig. 2d**). Some of the pathways commonly altered by eSTIM between humans and mice include cytokine activity, chemokine activity, chemokine receptor binding, cytokine receptor activity, and the CCR chemokine receptor binding. Some common genes enriched in these chemokine and cytokine-related pathways are *CCL3*, *CCL4*, *IL1A* and *TNF*. We further validated the expression of the IEG *Fos* and *Fosb*, and cytokine activity-related genes *Ccl3* and *Ccl4* using qPCR analysis from mouse brain following eSTIM (**Fig. 2e**). Thus, our findings of gene expression signatures in response to eSTIM in the human and mouse brains reveal a molecular signature that is partially conserved across species.

**Figure 2.**
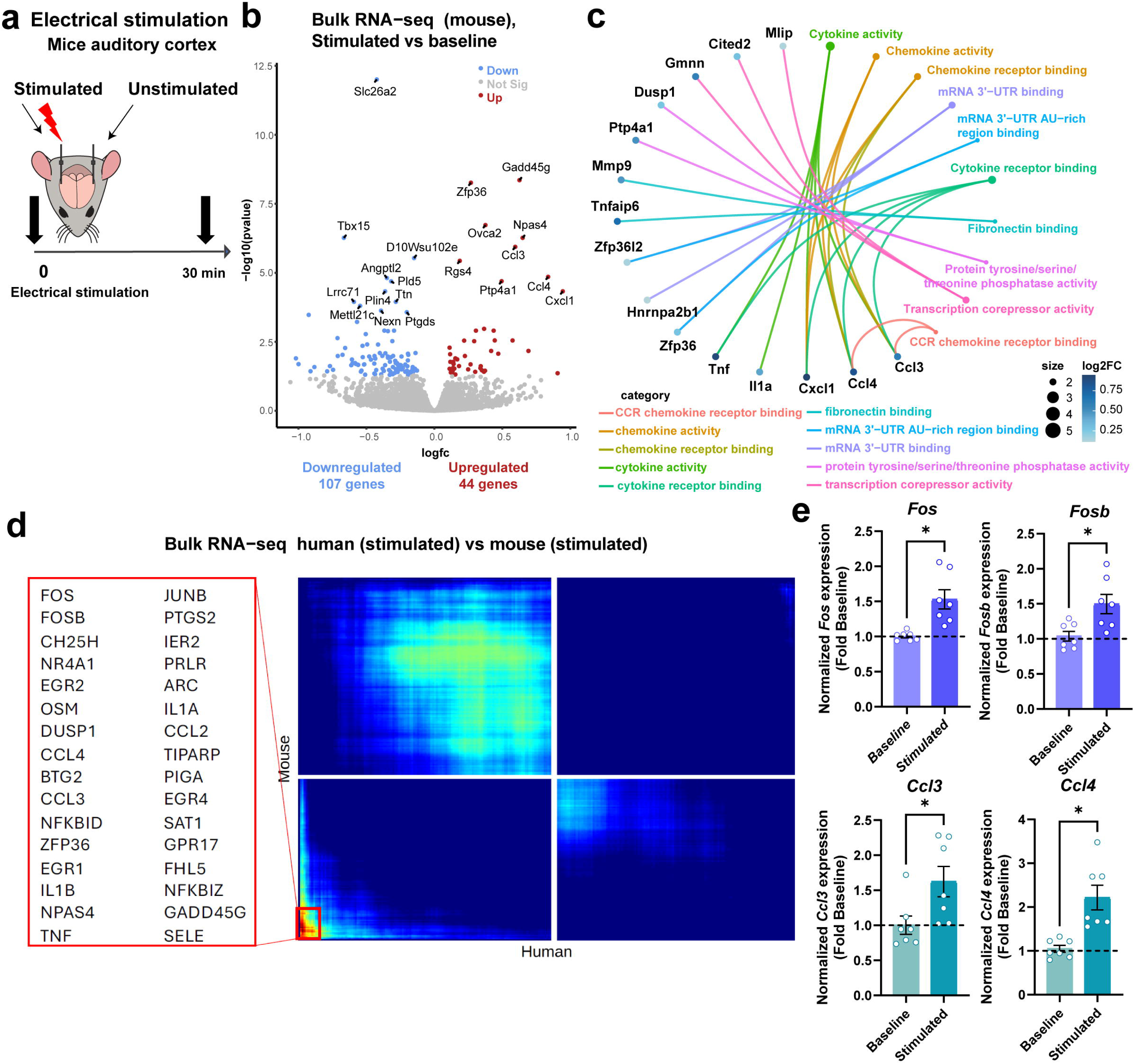
eSTIM of mouse cortex induces changes in genes associated with cytokine activity and transcription regulation. **a.** The mouse auditory cortex was stimulated, tissue was collected 30 min later, and the contralateral auditory cortex was obtained as baseline control. **b.** Volcano plot showing gene expression changes after stimulation. The most significant genes (FDR < 0.05) are labeled in red (upregulated) or blue (downregulated). **c.** Cnet plot showing pathway enrichment analysis of the genes significantly (FDR < 0.05) affected by eSTIM. **d.** Conserved gene expression signature across species. RRHO analysis demonstrates a significant overlap between genes upregulated in stimulated human samples and those from stimulated mouse samples. The heatmap shows the −log10(p-value) of the gene list overlap, with the bottom left red quadrant indicating significant co-upregulation. The list of 32 genes identified in this conserved signature is shown. **e.** qPCR analysis of the genes related to cytokine activity comparing stimulated versus baseline controls in mice cortex. Wilcoxon matched-pared signed rank test: Ccl3: p=0.0156, Ccl4:p=0.0156. n=7/group. Error bars represent ± SEM.

Next, we investigated the cell types exhibiting differential gene expression following eSTIM in the human brain by utilizing single-nuclei multi-omics (RNA and ATAC) on samples from the stimulated paradigm from four participants undergoing epilepsy resection surgery (**Fig 3a**). We also included samples from two participants with brain cancer undergoing glioblastoma resection surgery with the same stimulation parameters (**Fig 3a, Supplemental Table 1**). Cell clustering analysis identified six major cell types in our samples—excitatory neurons, microglia, inhibitory neurons, oligodendrocytes, oligodendrocytes precursor cells (OPC), and astrocytes (**Fig. 3b-c**). To determine the differentially expressed genes (DEGs) in each cell type following eSTIM, we performed a pseudobulk RNA-seq analysis due to its superior performance for detecting differential expression in single-cell RNA-sequencing analyses^33^ and the stringent nature of this analysis (**Supplementary Fig. 3**). The pseudo-bulk approach was more appropriate due to the high variability in the composition of cell types across samples (**Supplementary Fig. 4**). Mixed-effects statistical models were performed on all samples from glioblastoma and epilepsy patients together or in separate models (see Methods section; **Supplementary Fig. 5**). The pseudobulk analysis revealed 13 upregulated genes in excitatory neurons, 11 upregulated and 3 downregulated genes in inhibitory neurons (**Fig. 3d**). Classical IEGs such as *FOS*, *NPAS4*, and *EGR4* were upregulated in both excitatory and inhibitory neurons. Microglia displayed the highest differential gene expression of all cell types, with 135 upregulated and 8 downregulated genes (**Fig. 3d, Supplementary Fig. 5, Supplementary Table 5**). Genes related to cytokine activity (*CCL4L2*, and *IL1B*) were upregulated exclusively within microglia (**Supplementary Fig. 6**). Among the classical IEGs, eSTIM induced *FOS* expression in neurons (excitatory and inhibitory), microglia, oligodendrocytes, and OPC, while *FOSB*, *GADD45B* and *IER2* were upregulated in non-neuronal cells (microglia, oligodendrocytes, astrocytes and OPC) (**Supplementary Fig. 6**).

**Figure 3.**
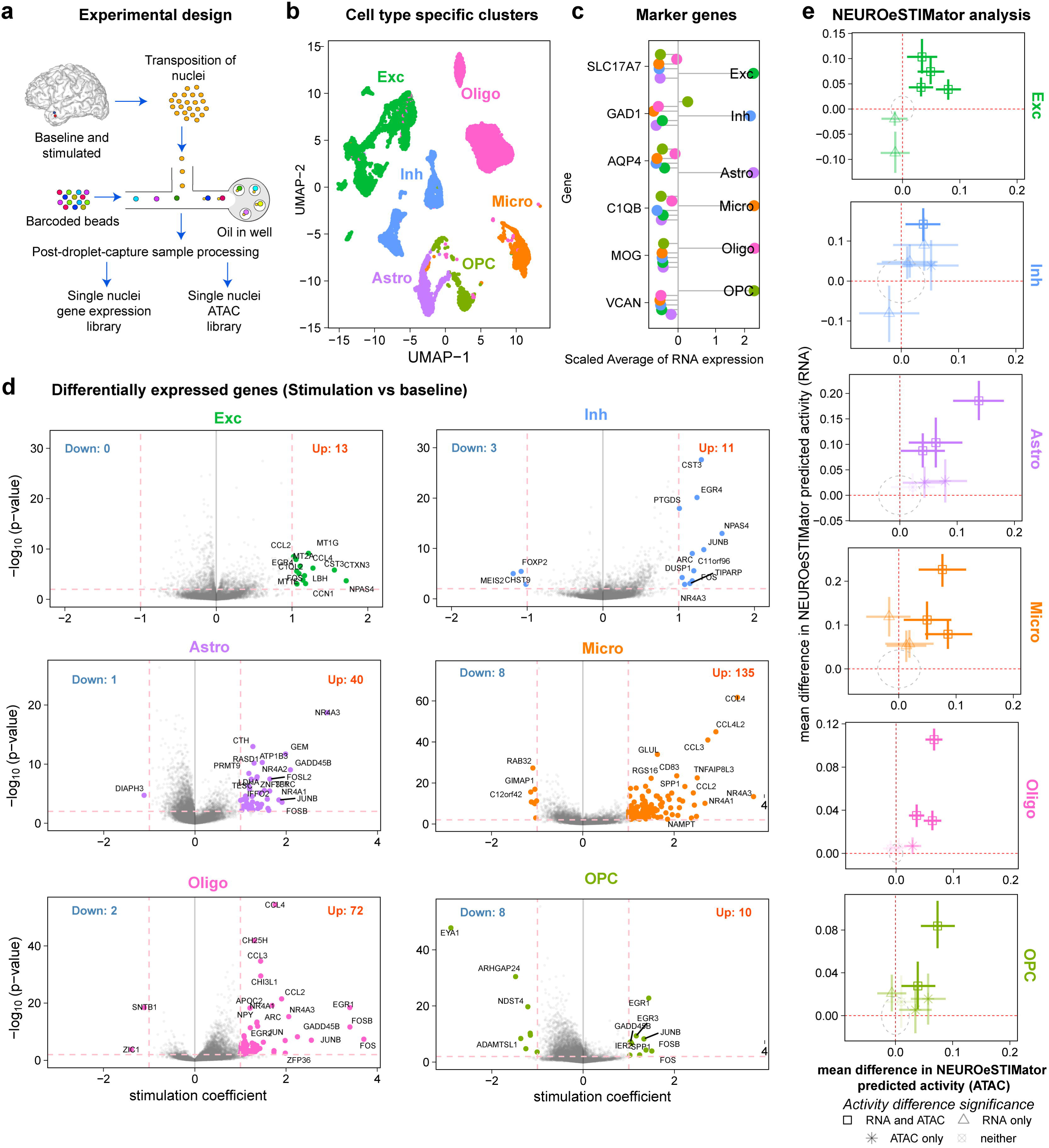
Single-nuclei multi-omics reveal cell type-specific transcriptomic changes following eSTIM in human cortex. **a**. Experimental scheme: Single-nucleus multi-omic experimental approach. **b.** UMAP shows the specific cell types from each cluster. **c.** Expression of key cell type markers in snRNA data. Values represent the mean expression across all samples for each cell type. **d.** Volcano plot showing differentially expressed genes in all cell types using pseudobulk analysis comparing stimulated vs baseline samples from the human brain. **e.** Scatter plots with error bars displaying bootstrapped mean differences in activity scores (stimulated - unstimulated) predicted by NEUROeSTIMator using ATAC-seq (x-axis) and RNA-seq (y-axis) inputs across major cell types. Each point represents the mean activity difference for an individual participant, derived from 1,000 bootstrap iterations. Error bars indicate 95% confidence intervals around the mean difference estimates. Point styling indicates statistical significance of activity differences: solid squares (both RNA and ATAC significant), triangles (RNA only significant), asterisk (ATAC only significant), and transparent squared-circles (neither significant). Dashed ellipses represent the null distribution generated by random shuffling of condition labels.

Comparing the upregulated genes in snRNA seq with human and mouse bulk RNA seq revealed 8 genes in microglia (*IL1A*, *TNF*, *FOS, FOSB*, *DUSP1*, *CCL3*, *CCL4*, and *ZFP36*) that are commonly upregulated following eSTIM in both groups (**Supplementary Fig. 7**). Other genes such as *CCL4L2*, *CD69*, *CD83*, *CDKN1A*, *IL1B*, *NEDD9*, *NFKBID*, *OLR1*, *OSM*, *RGS1*, *SOCS3*, and *TNFSF18* were identified as being specifically upregulated in human microglia after eSTIM, with no changes observed in mice (**Supplementary Figs. 7 and 8**). Gene ontology (GO) terms and pathways enriched among the upregulated genes in microglia included cytokine-related pathways such as cytokine activity, chemokine activity, CCR chemokine receptor, and cytokine receptor binding (**Supplementary Fig. 9, Supplementary Table 6-7**). Microglia exhibited increased expression of genes related to transcription regulatory pathways (such as DNA-binding transcription activator activity and cis-regulatory region sequence-specific DNA binding). These results reveal that eSTIM increases the expression of genes that are involved in cytokine activity and transcription regulation in microglia.

To further confirm that the differentially expressed genes identified by snRNA-seq (**Fig. 3d**) are specifically induced by eSTIM, rather than being affected by other factors such as disease conditions^34,35^ or neurosurgical procedures, we analyzed the overlap of DEGs from eSTIM with transcriptomic profiles of genes identified from studies from epilepsy^29,36^, glioblastoma^37^, memory oscillations^25,38^, and neurosurgery^39^ (**Supplementary Fig. 10**). Our results confirmed that cytokine-related genes *CCL4*, *CCL3*, and *CCL4L2* did not overlap with any transcriptomic profiles in the other datasets studied. However, our comparison with published data reveals overlap with epilepsy-associated genes in excitatory neurons^36^. Because our within-subject design includes baseline samples from the same participant, our findings reveal that the epilepsy-associated genes are specifically induced by eSTIM.

Next, we examined the cell type-specific response to eSTIM from within participants using our newly developed deep-learning computational tool, NEUROeSTIMator^40^, to map the active population of cells following eSTIM in each participant, comparing stimulated samples with corresponding baseline controls. NEUROeSTIMator, a deep-learning computational model was trained to infer neuronal activation from transcriptomic profiles and outputs a continuous activation score based on the expression of a curated set of genes known to be induced by neuronal activity and has been validated against electrophysiological properties using Patch-seq data, and is robust across species, cell types, and brain regions^40^. We estimated predicted activity using the NEUROeSTIMator tool on the snRNA-seq and snATAC-seq peaks using a window of ±2kb from the transcription start site (TSS) following eSTIM and corresponding baseline samples from each participant (**Fig. 3e**). We used bootstrap sampling to assess differences in predicted NEUROeSTIMator activity between stimulated and baseline for each cell type. Interestingly, we found that electrical stimulation significantly increased predicted activity seen from both RNA and ATAC samples in excitatory neurons in four out of six participants. In inhibitory neurons, we found that electrical stimulation significantly increased predicted activity (RNA and ATAC) in one out of six participants (**Fig. 3e)**. We also found that the microglia showed a significant increase in RNA and ATAC samples following eSTIM in three out of six participants, while all the six participants showed increased predicted activity from the RNA samples (**Fig. 3e**). These findings are consistent with our bulk RNA seq and snRNAseq data that eSTIM induces transcriptional signatures associated with transcriptional activation.

To identify the genome-wide transcription factor (TF) binding motifs that are enriched following eSTIM, we analyzed the snATAC-seq data and verified the cell identity by examining the accessibility of promoter regions for marker genes (**Fig. 4a-b**). Then, we examined genome-wide transcription factor binding motifs within the peaks that were upregulated after eSTIM across different cell types. The TF motif analysis revealed an enrichment of binding motifs of EGR sub-family members in multiple cell types (**Fig. 4c-d, Supplementary Fig. 11, and Supplementary Table 8**). Notably, the expression of several EGR sub-family members from snRNA-seq was also elevated following eSTIM (**Fig. 4c**). In microglia, motifs for EGR1, EGR2, and EGR3 were enriched after eSTIM, with the expression levels of these genes also increasing (**Fig. 4c, Supplementary Fig. 11**). Excitatory and inhibitory neurons showed induction of EGR4 gene expression and greater motif enrichment following eSTIM (**Fig. 4c, Supplementary Fig. 11**). Overall, our sn-ATAC-seq data reveal a signature of transcription factor motifs and chromatin accessibility that support gene expression changes caused by eSTIM in human microglia.

**Figure 4.**
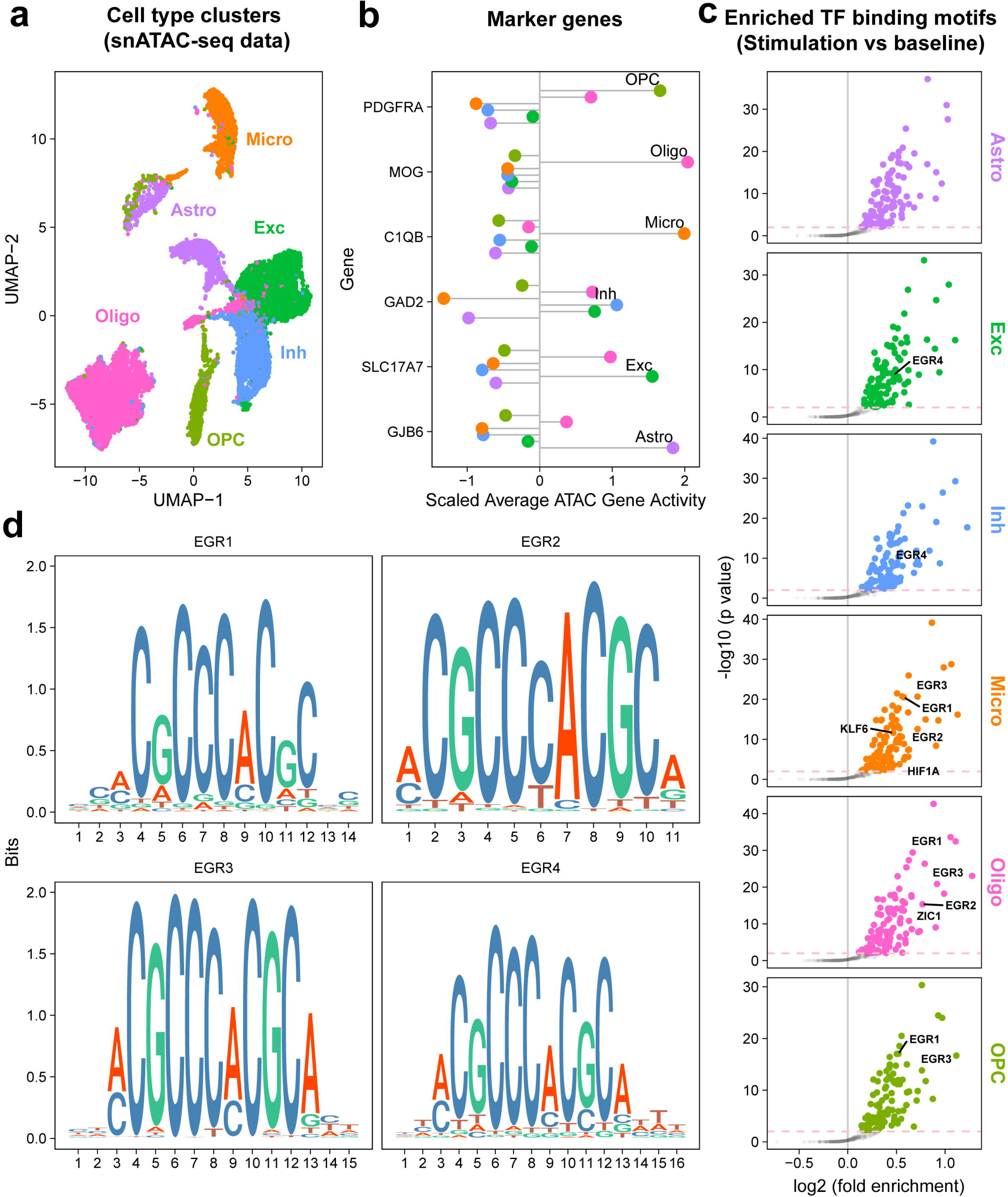
Chromatin accessibility changes mediated by eSTIM in human cortex. **a.** UMAP showing cell type cluster from the snATAC-seq data. **b.** Lolli-pop plot showing average gene activity changes due to chromatin accessibility changes on marker genes from the snATAC-seq data. **c.** Volcano plot showing significantly overrepresented DNA motifs in the set of peaks differentially accessible across different cell types in human cortex following eSTIM. Genes that also show significant changes in snRNA-seq data are labeled in the volcano plots. **d.** Motifs of TFs EGR1, EGR2, EGR3, and EGR4 are shown.

Neuropsychological assessment was available in 8 out of the 11 patients, showing that the patients did not have acquired global or focal deficits classically linked to the brain regions sampled (see Methods and Supplementary Table 1).

## Discussion

eSTIM is a model for studying the human cortex with protocols that are used clinically for mapping the function of specific brain regions. Here, we take a molecular approach, defining the transcriptomic and epigenomic signatures of eSTIM at the single-cell level in the human cortex in neurosurgical patient participants. Researchers have correlated human brain transcriptomics with prior recorded oscillatory signatures of memory consolidation^25,38^, but have not previously examined the changes in gene expression that are rapidly and directly driven by eSTIM *in vivo*. Our study is the first to demonstrate gene expression changes *in vivo* in the human brain caused by electrical stimulation, using a well-established clinical brain mapping protocol. By analyzing rare resection samples from living human cortex, we find that eSTIM impacts a network of cells that extends beyond excitatory neurons to include inhibitory neurons, astrocytes, oligodendrocytes and microglia.

Our single-nuclei transcriptomic data showed upregulation of IEGs *NPAS4*, *FOS* and *EGR4* in excitatory and inhibitory neurons. NPAS4 encodes a master transcription regulator of learning-induced genes and is critical for memory consolidation^41,42^. Studies in mice suggest that activity-induced expression of Npas4 mediates responses to adapt to homeostatic responses at a circuit-wide level^43^. Npas4 has a role in communicating increases in the spiking output and synaptic inputs of a neuron to tailor gene expression to the type of depolarizing activity^41,44^. Additionally, our findings that increased expression of classical IEGs such as *FOS*, *ARC*, *JUNB*, *EGR4*, and *DUSP1* in excitatory neurons suggest an early transcriptional response due to eSTIM. Transcription factors (TF) such as FOS, NPAS4, and EGR4 further regulate the expression of downstream gene targets, also known as late-response genes. Our study of chromatin accessibility defines a cell-type-specific signature of transcription factor binding motifs driving these processes that includes motifs for EGR1, EGR2, and EGR3, in microglia, and EGR4 in neurons. Previous work on rodent brains following electrical stimulation showed a long-lasting induction of *Egr1* (also known as *Zif268*), which correlates with the duration of long-term potentiation^45^. EGR1 induction and activation of microglia have also been reported *in vitro*^46^. Overall, our data suggests that eSTIM mediated molecular (transcriptomic and epigenomic) changes in the human brain may have a lasting effect on brain function.

Most work thus far has emphasized the role of neurons in responding to external stimuli, but we also found striking changes in non-neuronal cells, including microglia. In microglia, genes related to cytokine signaling showed the greatest induction pattern, and this was partially conserved across species (mouse and human). Microglia are critical modulators of neuronal function, acting to suppress excessive activity by inhibiting surrounding neurons, including excitatory neurons^47^. Although classified as non-excitable cells, microglia exhibit electrophysiological stimulus-response features, and changes in their membrane potential affect crucial microglial functions, including phagocytosis, chemotaxis, and cytokine release^48^. Our analyses identified critical molecular components within microglia that change with eSTIM, including chemokine-encoding genes such as *CCL3* and *CCL4*, which act to alter microglial motility, influence neuronal-microglial interactions, and shape neuronal connectivity^47,49–52^. These chemokines are ligands for C-C chemokine receptor type 5 (CCR5), which regulates neuronal excitability and memory allocation in the hippocampus^53^. A recent study investigating eSTIM induced genes in *ex vivo* human resection brain samples showed a similar signature of upregulation of these chemokine-encoding genes (*CCL3* and *CCL4*)^54^. Previous rodent studies have found that neuronal activation using chemogenetic approaches leads to upregulation of these genes (*CCL3* and *CCL4*) in microglia, including genes encoding chemokines^47^. Our work also revealed that genes encoding transcription factors such as FOS and NR4A1 are induced in microglia in response to eSTIM, thus suggesting a broad potential functional impact of these IEGs beyond their more frequently studied roles in neurons. Indeed, NR4A1 acts in neurons as a transcription factor essential for memory consolidation^55,56^ and functions in microglia as a molecular rheostat, contributing to the maintenance of a threshold that prevents microglial activation^57^. Therefore, as expected, we detected activity-dependent gene induction in neurons, however, identifying a microglial transcriptomic response following eSTIM represents an important conceptual advance in our understanding of the impact of this common clinically used protocol. Although our findings could not answer whether microglia directly respond to stimulation or to the changes in circuit activity induced by eSTIM, previous work suggest that microglia does respond directly following neuronal activation in mice^47^. Therefore, these findings suggest that at least a sub-set of microglial genes could respond due to circuit activity. Future work will be needed to resolve these questions and to determine whether neurons or non-neuronal cells are the primary drivers of the impact of eSTIM. Although these questions can be examined in mice or human brain slices, they would not address the fundamental question about how the human cortex is affected *in vivo,* which will require breakthroughs in FDA approved patient approaches.

This study represents a rare opportunity to define the activity-induced molecular changes in neurosurgical patients. In particular, the advance in our work is that we focus on the human brain and demonstrate that it is possible to study activity-induced transcriptomic changes *in vivo*. However, there are significant limitations to the experimental tools accessible to such studies. Although presenting a unique opportunity, it is important to note the caveats that come from this research carried out in a setting focused on clinical efficacy in patients that have epilepsy or glioblastoma. Thus, we cannot control all of the parameters as we might in an experiment carried out in a model organism, although we have worked to address three potential limitations of our study. First, our experimental design includes a group that does not receive eSTIM, and within subject baseline control, thus controlling for the impact of the disease, surgical procedure, craniotomy, brain temperature changes, and levels of anesthesia when present. We also show that eSTIM in a mouse model, in which we can control many additional parameters, gives rise to a partially conserved transcriptomic change. Importantly, our mouse experiment included the use of an inactive electrode, thus controlling for mechanically induced changes in gene expression. Second, the stimulation pattern used is safe and efficacious for brain mapping in human patients but is not optimized to modulate activity in specific circuits and cell types in the brain. Indeed, the stimulation protocol used leads to the functional disruption of brain regions^1^. Third, some of these patients have severe therapy-refractory epilepsy. However, the resected samples were taken a considerable distance from the seizure foci (see **Suppl. Fig. 1b**).

Despite these caveats, we were able to identify a reliable cell type-specific signature of the impact of eSTIM in the living human brain. This study demonstrates for the first time a unique transcriptomic signature in neurons and non-neuronal cells following direct eSTIM, representing a conceptual advance in our understanding of how the brain responds to an important clinical tool used to map brain function. These findings have the potential to inform the clinical practice of diagnostic and therapeutic brain stimulation. Clinical and translational research has focused mainly on the immediate effects of stimulation in neuronal populations, while microglia receive little attention, often being discussed with regard to neuroimmunological aspects^58^. Although for clinical mapping, clinicians rely on the immediate neuronal effects on behavior (e.g., speech arrest^10^), there is growing evidence that the therapeutic effects of eSTIM can accumulate gradually over time^59^. We could not test a range of stimulation frequencies and intensities or examine a broad range of brain areas and timepoints because of clinical limitations, but our results raise the intriguing hypothesis that microglia, not just neurons, shape plasticity after electrical stimulation. Our work further highlights that cytokine and chemokine-based mechanisms enable microglia to respond to eSTIM and possibly sculpt circuit function, making these potential targets to modify circuit activity with pharmacological approaches. Therefore, it will be especially interesting in future work to extend our studies to stimulation protocols used therapeutically, as well as natural stimuli to probe the critical role of neuronal and non-neuronal cells in mediating the response of the human brain to experience.

## Acknowledgments

We thank Rashmi N. Mueller who oversaw anesthesia to manage pain levels before, during, and after surgical procedures, and Kyle S. Conway for helping with the initial pathological assessment. We thank the Iowa Institute of Human Genetics (IIHG) core for RNA seq library preparation and sequencing. We thank Dr. Sourav Banerjee for providing valuable input on the manuscript.

## Funding

This work was supported by grants from the National Institute of Health R01 MH 087463 to T.A and The University of Iowa Hawkeye Intellectual and Developmental Disabilities Research Center (HAWK-IDDRC) P50 HD103556 to T.A., The National Institute of Health R00 AG068306 to S.C. and the National Institute of Health R01 DC004290 to M.A.H., National Science Foundation BCS-2342847 to C.I.P. T.N.J & C.I.P. are supported by a Research Program of Excellence of the Carver Trust and the Iowa Neuroscience Institute. K.Y. is supported by the University of Iowa CTSA K12 Program (K12TR004382). T.A. is supported by the Roy J. Carver Chair of Neuroscience, J.J.M is supported by the Roy J. Carver Professor of Neuroscience and the Andrew H. Woods Professorship supported T.N.J. The Neural Circuits and Behavior Core as well as the NeuroBank Core are supported by the Roy J. Carver Charitable Trust.

## Author contributions

S.C., M.A.H., and T.A. conceived the project, designed the experiments, and interpreted the data. S.C. and T.A. wrote the manuscript with assistance from C.I.P., L.L., and T.N.J., and inputs from other co-authors. H.K. and J.D.G performed human surgery and electrical stimulation and resected the tissue. A.E.R and C.K.K. processed with IRB and MRI imaging registration. H.O, M.A.H, T.N-J, and K.V.N provided advice on surgical procedures and electrical stimulation. M.M.H. provided a pathological assessment. M.B. provided neuropsychological assessment. L.C.L., K.V.N., B.J.P., C.I.P., and Y.A.A. collected the tissue and images during resection. S.C., B.J.P., L.C.L., U.M., L.L. performed human or mouse tissue samples experiments. C.I.P. and Y.A.A. performed distance to the seizure site analysis. U.M. and L.L. performed mouse brain imaging and data analysis. M.E., Y.V., A.I.G., and B.B. performed the bioinformatics analysis with inputs from E.B. and J.J.M. S.A.H. performed the mouse surgery and electrical stimulation experiments. K.V.N, C.K.K, Z.K. Y.K, C.I.P, M.M.H, J.J.M, E.B, U.M, and T.N.J. provided input on data analysis and interpretation. All authors discussed the results and commented on the manuscript.

## Competing interests

T.A. is a scientific advisor to Aditum Bio and Radius Health and serves on the scientific advisory board of Embark Neuro. The other authors declare no conflicting interests.

## Materials and Methods

### Patient participants

The study participants were 11 adult neurosurgical patients (8 female, 3 male, age 19-63 years old, median age 45.5±13.5 years old) with medically refractory epilepsy (9 patients) or glioblastoma (2 patients). The epilepsy patients were undergoing surgical resection of seizure foci following non-invasive electroencephalography (EEG) or invasive iEEG monitoring. All epileptic patients were diagnosed with intractable epilepsy and underwent ATL resection surgery for epilepsy of various etiologies **(Supplementary Fig. 1a, Supplementary Table 1).**

Two of the patient participants, had a dysembryoplastic neuroepithelial tumor. 6 out of 11 participants were required to be awake during the surgical resection, and the rest were anesthetized. Eight patients were non-smokers, two were former smokers (#G1, #E4) and one was smoker (#E2). Participant’s age, sex, surgery, and awake or sedative information were recorded (**Supplementary Table 1)**. All participants were native English speakers, 10 were right-handed, 1 was left-handed, and all had left language dominance as determined by Wada tests. All participants underwent audiometric evaluation before the study, and none were found to have hearing deficits or word recognition scores deemed sufficient to affect the findings presented in this study. The vision was self-reported as normal or corrected to normal with glasses or contact lenses. As determined by standard neuropsychological assessments, cognitive function was in the average range in all participants. Research protocols were approved by the University of Iowa Institutional Review Board (IRB 201910791, 201911084) and the National Institutes of Health. Written informed consent was obtained from all participants.

### Neuropsychological assessment

Neuropsychological data were obtained from the epilepsy patients during their Phase 1 neuropsychological outpatient examinations and were available for 8 of the 9 patients included in our sample. Cognitive test data are summarized in **Supplementary Table 1**, and as follows. Across these patients, performance, on a test commonly used by neuropsychologists to estimate pre-morbid baseline general cognitive ability (the Wide Range Achievement Test, Fourth Edition, Word Reading subtest), was strongly correlated with the Full-Scale Intelligence Quotient (FSIQ) from the Wechsler Adult Intelligence Scales – Fourth Edition (r = 0.674), and no participant demonstrated a large discrepancy between these two metrics. Additionally, no patient’s performance on the Boston Naming Test, which is sensitive to left hemisphere temporal lobe dysfunction, fell below the cutoff commonly used to identify frank impairment (i.e., a T score of 30, which corresponds to the 2nd percentile; **Suppl. Table 1**.). There was only one patient whose sample came from the right temporal lobe (#E5). That patient performed well within expectations on several tests sensitive to right hemisphere temporal lobe dysfunction, such as the Faces I and Faces II subtests from the Wechsler Memory Scales, Third Edition, which are often used to measure right temporal function in the setting of epilepsy. Altogether, these data suggest that the patients in our sample did not have an acquired deficit in cognition broadly speaking, nor a frank focal deficit classically, linked to the sampled tissue.

### Procedure

Surgery was performed awake or under general anesthesia. Standard craniotomy was performed by the same senior neurosurgeon in all epileptic patients to reach the epilepsy focus for resection, which involved the anterior and medial temporal lobe in all patients except one who had a temporal encephalocele and surrounding anterolateral temporal cortical focus. Before neurosurgical ATL resection, cortical tissue from the anterior temporal lobe was sampled by the neurosurgeon and handed over to the research team for analysis. Glioblastoma surgery resections were awake procedures and performed in the left temporal (G1) and frontal lobes (G2) by two senior neurosurgeons. Brain tissue over the most superficial presentation of the lesions was sampled before the tumor resection started.

The experimental condition was defined by an electrical “stimulation paradigm” and a control “no-stimulation paradigm**”**. The location of the sampled tissue is plotted on anatomic brain reconstructions **(Supplementary Fig. 1b).** The 30 minutes in between the control and experimental samples was clinically required for the clinical EEG team to record inter-ictal activity with surface recording grids placed gently on the brain surface.

In the “stimulation paradigm”, a baseline sample was obtained from the anterior temporal cortex that would be resected in the planned surgical resection. The area directly adjacent to where the baseline sample was collected was stimulated with direct bipolar electric stimulation (50 Hz frequency, 0.2 ms pulse duration, 2 min stimulation duration, and 10 V voltage). The stimulated area was then sampled after a period of 30 minutes to allow for gene expression^60,61^. In the “no stimulation paradigm”, no direct electric stimulation was performed and the area directly adjacent to the baseline sample was collected 30 minutes after initial baseline sampling.

After sampling the tissue samples were immediately placed in a sterile container on dry ice. The average weight of the baseline sample was 88.0 ± 33.2 mg (mean, standard deviation) and the adjacent sample was 113.4 ± 62.7 mg. After the collection of all samples in this fashion, they were weighed and transferred to a freezer at −80° C for storage until further testing.

### Sample Localization to MNI space

Intraoperative photos of the sample sites were obtained during the time of surgery. Using patient participant matched preoperative T1 sequence MRI, the sample sites were mapped onto their anatomic brain reconstructions **(Supplementary Fig. 1a)**. They were also mapped onto MNI space coordinates.

### Human cortical tissue RNA extraction, library preparation, and sequencing

Total RNA was extracted from sampled human brains using miRNeasy Mini Kit (Qiagen, CA, USA). The tissue samples were homogenized in QIAzol (Qiagen, CA. USA) stainless steel beads (Qiagen, CA, USA). Chloroform was then used for phase separation. RNA containing an aqueous layer was further purified using the RNeasy MinElute spin column. RNA was finally eluted in RNase-free water. RNA concentrations were estimated using a Nanodrop (Thermo Fisher Scientific, MA, USA) and Qubit (Thermo Fisher Scientific, MA, USA). RNA libraries were prepared at the Iowa Institute of Human Genetics (IIHG), Genomics Division, using the Illumina Stranded Total RNA Prep, Ligation with Ribo-Zero Plus (Illumina Inc., San Diego, CA). The KAPA Illumina Library Quantification Kit (KAPA Biosystems, Wilmington, MA) was used to measure library concentrations. Pooled libraries were sequenced on Illumina NovaSeq6000 sequencers with 150-bp paired-end chemistry (Illumina) at the Iowa Institute of Human Genetics (IIHG) core.

### Bulk RNA sequencing analysis

RNA-seq data were processed with the bcbio-nextgen pipeline (https://github.com/bcbio/bcbio-nextgen, version 1.2.9). The pipeline uses STAR^62^ to align reads to the hg38 or mm10 reference genome and quantifies expression at the gene level with feature Counts ^63^. All further analyses were performed using R. For gene-level count data, the R package EDASeq was used to account for sequencing depth (upper quartile normalization) ^64^. Latent sources of variation in expression levels were assessed and accounted for using RUVSeq (RUVs mode using all features)^65^. Appropriate choice of the RUVSeq parameter k was guided through inspection of principal components analysis (PCA) plots. Specifically, the smallest value k was chosen where PCA plots demonstrated replicate sample clustering in the first three principal components^66^. Differential expression analysis was conducted using the edgeR package^67^. Codes to reproduce the RNA-seq differential gene expression analysis are available at https://github.com/YannVRB/Human-brain-stimulation.git.

All the transcriptomics data have been deposited in NCBI’s Gene Expression Omnibus and are accessible through GEO Series accession number GSE224952.

### Downstream pathway analysis

The enrichment analysis of differentially expressed genes-associated pathways and molecular functions from the stimulated and unstimulated RNA-seq was performed with the Gene Ontology (GO–molecular function) databases using clusterProfiler package in R. Only the pathways with an adjusted p-value < 0.05 were considered as significant and displayed. Further, the enrichment data were visualized using ‘cnetplot’ function of clusterProfiler.

#### Cross-species comparison using RRHO

To identify conserved transcriptional changes between human and mouse models, we performed a Rank-Rank Hypergeometric Overlap (RRHO) analysis ^31,32^. Differential expression results from the human stimulated paradigm **(Supplementary Table 2)** were compared to those from the mouse stimulated experiment **(Supplementary Table 4)**. For each species, genes were ranked by a signed −log10(p-value) score, which incorporates both statistical significance and the direction of fold change. The RRHO analysis was performed using the RRHO2 package in R (v.1.0.), with a step size of 50 and multiple testing correction using the Benjamini-Yekutieli method. Genes in the significant overlap regions were extracted, identifying conserved upregulated and downregulated gene sets between species.

#### Single-nuclei multiomics (nuclei isolation, library preparation, sequencing

Nuclei were isolated from brain tissue using the Chromium Nuclei Isolation Kit (10X Genomics). Briefly, frozen tissue was dissociated with pestle in lysis buffer, passed through nuclei isolation column and spun at 16,000 rcf for 20 sec at 4^0^C. Flowthrough was vortexed and spun at 500 rcf for 3 mins at 4^0^C. Pellet was resuspended with debris removal buffer and centrifuged at 700 rcf for 10 mins at 4^0^C, nuclei resuspended in wash buffer and centrifuged again at 500 rcf for 5 mins at 4^0^C. Pellet was resuspended in resuspension buffer and nuclei were counted using a hemocytometer. Nuclei were directly processed for droplet capture for single cell multiome ATAC + gene expression using a chromium controller (10X Genomics). Chromium Next GEM Single Cell Multiome ATAC + Gene v1 chemistry was used to create single nuclei ATAC and RNA libraries from the same cell. Libraries were sequenced on an Illumina Novaseq 6000 with a 150 bp paired end read setup.

### Single-nuclei multi-omics data processing

The single-nuclei multi-omics data from the human brain stimulation samples was processed to generate an RNA gene counts matrix and ATAC fragments using the Cell Ranger ARC (https://www.10xgenomics.com/support/software/cell-ranger-arc/latest) 2.0.2 pipeline from 10x Genomics. Each sample was processed and used to create a Seurat object in R, using the Seurat package ^68–72^. The initial quality control (QC) steps for the cells included filtering based on total count of ATAC peaks (> 1000 and < 100000), nucleosome signal (< 2), TSS enrichment (> 1), total count of RNA (> 1000 and < 40000), and percent of mitochondrial genes (< 20). (See **Supplementary Fig. 2** for the methods outline.)

For the ATAC data processing, we used the EnsDb.Hsapiens.v86 ^73^ as the main annotation for the identified fragments. We additionally called the ATAC peaks using the MACS3 ^74^ CallPeaks function instead of relying on the Cell Ranger ARC called peaks, as it merges distinct peaks that are close together, causing a problem for motif enrichment analysis and peak-to-gene linkage. The identified peaks were then filtered to keep peaks on standard chromosomes and drop genomic blacklist regions.

The RNA counts data were normalized per sample using the NormalizeData function from Seurat with the log normalization as the normalization method with other default function parameters. The normalized data was then scaled, and the top 2000 variable features were identified using vst as the selection method.

All Seurat samples were merged into one Seurat object to be used for the cell type annotation and dimensionality reduction methods. We used the Allen Institute dataset for human brain cell types ^75^ as a reference in cell type identification. The cell type labeling and predictions were done using the functions FindTransferAnchors and TransferData. In the cell type labeling step, we decided to proceed with the major cell type labeling instead of the refined subclasses per cell type.

In our second QC step for each identified cell type, we dropped cells that had a probability < 0.75 for the predicted cell type. Based on this criterion, certain cell types were completely dropped from certain samples, so we decided to not proceed with these cell types for downstream analysis. The dropped cell types included endothelial and VLMC cells. Cells that passed the mentioned criteria were then used for the pseudo-bulk step.

To overcome the dropout problem of the RNA counts from the data, we used DCA ^76^, a deep count autoencoder for denoising scRNA-seq data. Prior to the imputation step, we dropped genes with a total sum of 0 counts across all cells. DCA imputation was done using the raw gene counts from all samples with default parameters from the model, and the imputed values were then used for the remaining analysis. We did the imputation for baseline cells independently from the stimulated cells to avoid the potential bias in imputation based on the number of cells per condition.

Due to the high variability in samples’ composition (i.e., proportions of detected cell types in each sample; **Supplementary Fig. 4**), we used a sampling with replacement approach prior to the pseudo-bulk step. We sampled X number of cells from each cell type in each sample for 100 iterations. In each iteration, those sampled cells were aggregated to provide a pseudo-bulk sample. Then, the resulting 100 pseudo-bulk samples from each cell type per sample were averaged. The X number of sampled cells for aggregation was different per cell type as we sampled the rounded value of the 10^th^ percentile of cell count per cell type (Table 1).

**Table 1:** Rounded tenth percentile of cell count per cell type. These numbers were used for sampling with replacement and bootstrapping.

|  | Astro | Micro | Oligo | OPC | Exc | Inh |
| --- | --- | --- | --- | --- | --- | --- |
| 10<sup>th</sup> percentile | 46 | 41 | 380 | 92 | 112 | 32 |

The differential gene expression (DGE) analysis was performed using two different bulk RNA-Seq packages: edgeR ^67^ and the lmmSeq model from glmmSeq ^77^. Both packages use different models to estimate the coefficient for the variable of interest. We focused on lmmSeq in our main results as it relies on a mixed-effects model and considers the random effect of participants, aside from the main effect of stimulation. For each method, we had three different models: two by-dx models and a model for all samples combined. The by-dx models are independent models for each dx group (epilepsy/glioblastoma). We only show the results of the combined model in the main figures, and the remaining results are in **Supplementary Fig. 5**. The formulas for all models were as below:

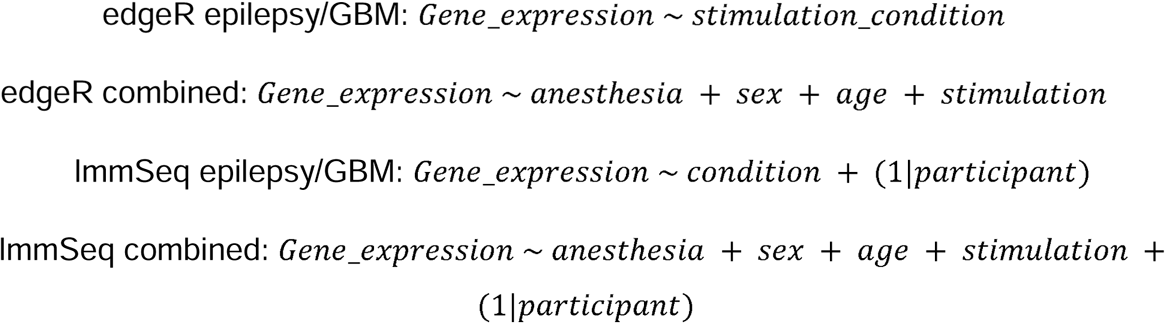

We used a binary representation for the stimulation condition (1=stimulated; 0=baseline), sex (1=male; 0=female), and anesthesia (1=present; 0=no anesthesia/awake) coefficients in the model. The dx variable was used as a factor variable with two levels: epilepsy and glioblastoma. The participant variable was used as a random variable in the mixed-effects model as a factor of 6 levels (i.e., 6 patients). Age was used as a numeric variable representing age in years. The epilepsy models were applied to the data from 4 patients (8 samples; each patient had both baseline and stimulated conditions).

We did not include covariates like sex, age, and anesthesia in any of the by-dx models because of the limited sample size per dx and the strong collinearity between parameters in the model that leads to estimation failure. We identified genes as being differentially expressed by using a threshold on both the coefficient for the stimulation parameter in the formula (> 1 or < −1) and the FDR (< 0.05) values. All models from both methods were tested independently for each cell type, and the results were compared.

### Bulk RNA-Seq deconvolution

We deconvolved the bulk RNA-Seq human samples using the granulator package in R with the Allen Institute cell types data set as a reference. We show that the cell type proportions in both stimulated and baseline conditions were not significantly different per participant (**Supplementary Fig. 2).**

### NEUROeSTIMator

To estimate the cell activity, we used NEUROeSTIMator and predicted the activity score (0-1) for each cell. The activity was estimated on the RNA-Seq level, and the gene activity was linked to peaks from the ATAC data. Then, we sampled the same X number of cells per cell type used for pseudo-bulk analysis for 1000 iterations, and estimated the mean with a 95% confidence interval for each sample. Finally, we investigated the difference between stimulated and baseline conditions per subject on a cell type level.

### Comparison to literature

To assess if the genes induced after electrical stimulation from the human neurosurgery are solely due to stimulation, we compared the identified DEGs to genes previously linked to epilepsy, glioblastoma, neurosurgery, and memory. For the epilepsy genes, we considered genes from the literature ^36^, differential expression from bulk and single-cell RNA-Seq ^29,36^, and a Genome-Wide Association Study (GWAS) ^78^. We also included DEGs from single-cell multi-omics data for glioblastoma ^37^ and oscillatory signatures supporting human episodic memory encoding ^25,38^. Additionally, we considered genes identified by the Allen Institute to be differentially expressed due to the neurosurgery ^39^, even though our protocol considers that effect by doing a within-sample design and having the human neurosurgery sample that did not have the stimulation paradigm applied (**Supplementary Fig. 10**).

### Enrichment Analysis

We performed gene ontology (GO) analysis for the identified DEGs from the mixed-effects model for each cell type using the goseq v.1.50.0 package in R ^79^. All GO terms were filtered based on their FDR to be < 0.05 and reported in **Supplementary Table 6**. Additionally, we used the ActivePathways R package ^80^ to check for enriched pathways based on the Human Reactome April 2025 release ^81^. Pathway enrichment was done using genes with FDR < 0.1 ranked based on their FDR value for each cell type.

#### Animals

Adult male C57BL/6J mice were purchased from The Jackson Laboratory were 3 to 4 months of age during experiments. All mice had free access to food and water; lights were maintained on a 12-hour light/12-hour dark cycle.

#### Mouse electrical stimulation

Stimulation experiments were performed in anesthetized adult male C57BL6/J mice. Anesthesia was induced with 5% isoflurane by inhalation and maintained at 1.8-2% for the duration of the experiment. The mouse was placed in a stereotax (Kopf) and a midline incision was made and the skin retracted to expose the temporal muscle bilaterally. The dorsal insertion of both temporal muscles was removed, and the muscles retracted. A 2-3 mm craniotomy was made over area AuV (centered at 2.9 mm posterior, 4.2 mm lateral, 2.8 mm ventral from Bregma based on Paxinos atlas) bilaterally to expose the cortical surface and a small square of gel foam soaked in ACSF was applied on top of the dura to prevent it from dehydrating. For each mouse the side of electrical stimulation and sham stimulation were alternated, with the sham side serving as the baseline control for gene expression profiling. Electrical stimulation was delivered through a bipolar ball electrode constructed from two silver wires in which the uninsulated tips were melted under a butane flame (1 mm tip size, 2 mm tip spacing). For both electrical and sham stimulations, the gel foam was removed and the electrode was gently lowered to make contact with the cortical surface. For electrical stimulation, a biphasic pulse train was then delivered for two minutes (8 mA, 50 Hz, 200 µs pulse width). For the sham stimulation no current was delivered but the electrode was left in place for 2 minutes. Following electrical or sham stimulation the electrode was slowly retracted, and the exposed dura was covered with gel foam. Selection for hemisphere was performed randomly, which resulted in the order of electrical and sham stimulation alternating from mouse to mouse. No more than 5 minutes elapsed between electrical or sham stimulation of both sides. After both sides were stimulated (electrical or sham) the mouse was left in the stereotax under anesthesia for 30 minutes before euthanasia and tissue collection. Tissue samples were immediately stored at −80^0^C in RNA later solution (Ambion).

#### Bulk RNA extraction, cDNA preparation and qPCRs from mouse auditory cortex

Tissue samples were homogenized in Qiazol (Qiagen) using stainless steel beads (Qiagen). Chloroform was added and centrifuged at 12,000*g* at room temperature for 15 min to separate RNA in the aqueous phase. RNA was precipitated in ethanol and cleared using RNeasy kit (Qiagen). RNA eluted in nuclease-free water was then treated with DNase (Qiagen) at room temperature for 25 min to remove genomic DNA. RNA was further precipitated in ethanol, sodium acetate (pH 5.2) and glycogen overnight at −20°C. RNA was precipitated by centrifugation at 30,000g for 20 min, precipitate washed with 70% ethanol and the dried RNA pellet was resuspended in nuclease-free water. RNA concentration was measured using NanoDrop (Thermo Fisher Scientific). 1 μg of RNA was used for complementary DNA (cDNA) preparation using the SuperScript IV First-Strand Synthesis System (Ambion). Real-time reverse transcription polymerase chain reactions (RT-PCRs) were performed on the QuantStudio 7 Flex Real-Time PCR System (Applied Biosystems, Life Technologies). Data were normalized to housekeeping genes (*Tubulin*, *Pgk1*, and *Actin*), and 2^(−ΔΔCt)^ method was used for gene expression analysis.

#### Statistics

For the qPCR analysis, the Wilcoxon matched pairs signed rank test and one sample Wilcoxon test was performed.

## Code availability statement

All codes for bulk RNA-Seq differential expression, analyzing single-nuclei mutli-omics (pre-processing, DCA imputation, pseudobulk, differential expression, enrichment analysis, and comparison to literature), running NEUROeSTIMator, and bulk RNA-Seq deconvolution can be found at https://github.com/melsadany/The-gene-expression-signature-of-electrical-stimulation-in-the-human-brain.

## Data availability statement

Bulk RNA-seq and single-nuclei multi-omics data, has been deposited in NCBI’s Gene Expression Omnibus and are accessible through GEO Series accession number GSE224952.

## Notes

### Competing Interest Statement

T.A. is a scientific advisor to Aditum Bio and Radius Health and he serves on the scientific advisory board of Embark Neuro. T.A serves on the Scientific Advisory Board of EmbarkNeuro and is a scientific advisor to Aditum Bio and Radius Health. The other authors declare no conflicting interests.

### Summary of Updates

Added a larger sample size that includes cancer patients in addition to epilepsy patients. Figure 3 and 4 revised with new computational analysis.

## References

1 Penfield, W. & Rasmussen, T. Vocalization and arrest of speech. Arch Neurol Psychiatry 61, 21–27 (1949). 10.1001/archneurpsyc.1949.02310070027002

2 Lozano, A. M. et al. Deep brain stimulation: current challenges and future directions. Nat Rev Neurol 15, 148–160 (2019). 10.1038/s41582-018-0128-2

3 Okun, M. S. Deep-brain stimulation for Parkinson’s disease. N Engl J Med 367, 1529–1538 (2012). 10.1056/NEJMct1208070

4 De Risio, L. et al. Recovering from depression with repetitive transcranial magnetic stimulation (rTMS): a systematic review and meta-analysis of preclinical studies. Transl Psychiatry 10, 393 (2020). 10.1038/s41398-020-01055-2

5 Kellner, C. H. et al. ECT in treatment-resistant depression. Am J Psychiatry 169, 1238–1244 (2012). 10.1176/appi.ajp.2012.12050648

6 Krauss, J. K. et al. Technology of deep brain stimulation: current status and future directions. Nat Rev Neurol 17, 75–87 (2021). 10.1038/s41582-020-00426-z

7 Sheth, S. A. & Mayberg, H. S. Deep Brain Stimulation for Obsessive-Compulsive Disorder and Depression. Annu Rev Neurosci 46, 341–358 (2023). 10.1146/annurev-neuro-110122-110434

8 Goodman, W. K., Storch, E. A. & Sheth, S. A. Harmonizing the Neurobiology and Treatment of Obsessive-Compulsive Disorder. Am J Psychiatry 178, 17–29 (2021). 10.1176/appi.ajp.2020.20111601

9 Kombos, T. & Suss, O. Neurophysiological basis of direct cortical stimulation and applied neuroanatomy of the motor cortex: a review. Neurosurg Focus 27, E3 (2009). 10.3171/2009.8.FOCUS09141

10 Lu, J. et al. Functional maps of direct electrical stimulation-induced speech arrest and anomia: a multicentre retrospective study. Brain 144, 2541–2553 (2021). 10.1093/brain/awab125

11 George, D. D., Ojemann, S. G., Drees, C. & Thompson, J. A. Stimulation Mapping Using Stereoelectroencephalography: Current and Future Directions. Front Neurol 11, 320 (2020). 10.3389/fneur.2020.00320

12 Grande, K. M., Ihnen, S. K. Z. & Arya, R. Electrical Stimulation Mapping of Brain Function: A Comparison of Subdural Electrodes and Stereo-EEG. Front Hum Neurosci 14, 611291 (2020). 10.3389/fnhum.2020.611291

13 Puk, O., Jablonska, M. & Sokal, P. Immunomodulatory and endocrine effects of deep brain stimulation and spinal cord stimulation - A systematic review. Biomed Pharmacother 168, 115732 (2023). 10.1016/j.biopha.2023.115732

14 Yap, E. L. & Greenberg, M. E. Activity-Regulated Transcription: Bridging the Gap between Neural Activity and Behavior. Neuron 100, 330–348 (2018). 10.1016/j.neuron.2018.10.013

15 Sagar, S. M., Sharp, F. R. & Curran, T. Expression of c-fos protein in brain: metabolic mapping at the cellular level. Science 240, 1328–1331 (1988). 10.1126/science.3131879

16 Guzowski, J. F., McNaughton, B. L., Barnes, C. A. & Worley, P. F. Environment-specific expression of the immediate-early gene Arc in hippocampal neuronal ensembles. Nat Neurosci 2, 1120–1124 (1999). 10.1038/16046

17 Laricchiuta, D. et al. Optogenetic Stimulation of Prelimbic Pyramidal Neurons Maintains Fear Memories and Modulates Amygdala Pyramidal Neuron Transcriptome. Int J Mol Sci 22 (2021). 10.3390/ijms22020810

18 Bali, P. & Kenny, P. J. Transcriptional mechanisms of drug addiction Dialogues Clin Neurosci 21, 379–387 (2019). 10.31887/DCNS.2019.21.4/pkenny

19 Fernandez-Albert, J. et al. Immediate and deferred epigenomic signatures of in vivo neuronal activation in mouse hippocampus. Nat Neurosci 22, 1718–1730 (2019). 10.1038/s41593-019-0476-2

20 Marco, A. et al. Mapping the epigenomic and transcriptomic interplay during memory formation and recall in the hippocampal engram ensemble. Nat Neurosci 23, 1606–1617 (2020). 10.1038/s41593-020-00717-0

21 Trevino, A. E. et al. Chromatin and gene-regulatory dynamics of the developing human cerebral cortex at single-cell resolution. Cell 184, 5053–5069 e5023 (2021). 10.1016/j.cell.2021.07.039

22 Reijmers, L. G., Perkins, B. L., Matsuo, N. & Mayford, M. Localization of a stable neural correlate of associative memory. Science 317, 1230–1233 (2007). 10.1126/science.1143839

23 Halder, R. et al. DNA methylation changes in plasticity genes accompany the formation and maintenance of memory. Nat Neurosci 19, 102–110 (2016). 10.1038/nn.4194

24 Boulting, G. L. et al. Activity-dependent regulome of human GABAergic neurons reveals new patterns of gene regulation and neurological disease heritability. Nat Neurosci 24, 437–448 (2021). 10.1038/s41593-020-00786-1

25 Berto, S. et al. Gene-expression correlates of the oscillatory signatures supporting human episodic memory encoding. Nat Neurosci 24, 554–564 (2021). 10.1038/s41593-021-00803-x

26 Konopka, G. Cognitive genomics: Linking genes to behavior in the human brain. Netw Neurosci 1, 3–13 (2017). 10.1162/NETN_a_00003

27 Berger, M. S. & Ojemann, G. A. Intraoperative brain mapping techniques in neuro-oncology. Stereotact Funct Neurosurg 58, 153–161 (1992). 10.1159/000098989

28 Sanai, N., Mirzadeh, Z. & Berger, M. S. Functional outcome after language mapping for glioma resection. N Engl J Med 358, 18–27 (2008). 10.1056/NEJMoa067819

29 Brueggeman, L. et al. Drug repositioning in epilepsy reveals novel antiseizure candidates. Ann Clin Transl Neurol 6, 295–309 (2019). 10.1002/acn3.703

30 Altmann, A. et al. A systems-level analysis highlights microglial activation as a modifying factor in common epilepsies. Neuropathol Appl Neurobiol 48, e12758 (2022). 10.1111/nan.12758

31 Cahill, K. M., Huo, Z., Tseng, G. C., Logan, R. W. & Seney, M. L. Improved identification of concordant and discordant gene expression signatures using an updated rank-rank hypergeometric overlap approach. Sci Rep 8, 9588 (2018). 10.1038/s41598-018-27903-2

32 Plaisier, S. B., Taschereau, R., Wong, J. A. & Graeber, T. G. Rank-rank hypergeometric overlap: identification of statistically significant overlap between gene-expression signatures. Nucleic Acids Res 38, e169 (2010). 10.1093/nar/gkq636

33 Murphy, A. E. & Skene, N. G. A balanced measure shows superior performance of pseudobulk methods in single-cell RNA-sequencing analysis. Nat Commun 13, 7851 (2022). 10.1038/s41467-022-35519-4

34 Vezzani, A., French, J., Bartfai, T. & Baram, T. Z. The role of inflammation in epilepsy. Nat Rev Neurol 7, 31–40 (2011). 10.1038/nrneurol.2010.178

35 Devinsky, O., Vezzani, A., Najjar, S., De Lanerolle, N. C. & Rogawski, M. A. Glia and epilepsy: excitability and inflammation. Trends Neurosci 36, 174–184 (2013). 10.1016/j.tins.2012.11.008

36 Pfisterer, U. et al. Identification of epilepsy-associated neuronal subtypes and gene expression underlying epileptogenesis. Nature Communications 11, 5038 (2020). 10.1038/s41467-020-18752-7

37 Wang, X. et al. Single-cell multi-omics sequencing uncovers region-specific plasticity of glioblastoma for complementary therapeutic targeting. Sci Adv 10, eadn4306 (2024). 10.1126/sciadv.adn4306

38 Berto, S., Wang, G. Z., Germi, J., Lega, B. C. & Konopka, G. Human Genomic Signatures of Brain Oscillations During Memory Encoding. Cereb Cortex 28, 1733–1748 (2018). 10.1093/cercor/bhx083

39 Hodge, R. D. et al. Conserved cell types with divergent features in human versus mouse cortex. Nature 573, 61–68 (2019). 10.1038/s41586-019-1506-7

40 Bahl, E. et al. Using deep learning to quantify neuronal activation from single-cell and spatial transcriptomic data. Nat Commun 15, 779 (2024). 10.1038/s41467-023-44503-5

41 Fu, J., Guo, O., Zhen, Z. & Zhen, J. Essential Functions of the Transcription Factor Npas4 in Neural Circuit Development, Plasticity, and Diseases. Front Neurosci 14, 603373 (2020). 10.3389/fnins.2020.603373

42 Ramamoorthi, K. et al. Npas4 regulates a transcriptional program in CA3 required for contextual memory formation. Science 334, 1669–1675 (2011). 10.1126/science.1208049

43 Spiegel, I. et al. Npas4 regulates excitatory-inhibitory balance within neural circuits through cell-type-specific gene programs. Cell 157, 1216–1229 (2014). 10.1016/j.cell.2014.03.058

44 Brigidi, G. S. et al. Genomic Decoding of Neuronal Depolarization by Stimulus-Specific NPAS4 Heterodimers. Cell 179, 373–391 e327 (2019). 10.1016/j.cell.2019.09.004

45 Abraham, W. C. et al. Correlations between immediate early gene induction and the persistence of long-term potentiation. Neuroscience 56, 717–727 (1993). 10.1016/0306-4522(93)90369-q

46 Langmann, T. et al. Induction of early growth response-1 mediates microglia activation in vitro but is dispensable in vivo. Neuromolecular Med 11, 87–96 (2009). 10.1007/s12017-009-8061-6

47 Badimon, A. et al. Negative feedback control of neuronal activity by microglia. Nature 586, 417–423 (2020). 10.1038/s41586-020-2777-8

48 Laprell, L., Schulze, C., Brehme, M. L. & Oertner, T. G. The role of microglia membrane potential in chemotaxis. J Neuroinflammation 18, 21 (2021). 10.1186/s12974-020-02048-0

49 Liu, Y. U. et al. Neuronal network activity controls microglial process surveillance in awake mice via norepinephrine signaling. Nat Neurosci 22, 1771–1781 (2019). 10.1038/s41593-019-0511-3

50 Stowell, R. D. et al. Noradrenergic signaling in the wakeful state inhibits microglial surveillance and synaptic plasticity in the mouse visual cortex. Nat Neurosci 22, 1782–1792 (2019). 10.1038/s41593-019-0514-0

51 Cheadle, L. et al. Sensory Experience Engages Microglia to Shape Neural Connectivity through a Non-Phagocytic Mechanism. Neuron 108, 451–468 e459 (2020). 10.1016/j.neuron.2020.08.002

52 Estevao, C. et al. CCL4 induces inflammatory signalling and barrier disruption in the neurovascular endothelium. Brain Behav Immun Health 18, 100370 (2021). 10.1016/j.bbih.2021.100370

53 Shen, Y. et al. CCR5 closes the temporal window for memory linking. Nature 606, 146–152 (2022). 10.1038/s41586-022-04783-1

54 Moore, H. et al. Stimulation modulates cell assemblies linked with gene networks in the human temporal cortex ex vivo. bioRxiv (2025). 10.1101/2025.06.25.661589

55 Chatterjee, S. et al. Endoplasmic reticulum chaperone genes encode effectors of long-term memory. Sci Adv 8, eabm6063 (2022). 10.1126/sciadv.abm6063

56 Vanrobaeys, Y. et al. Mapping the spatial transcriptomic signature of the hippocampus during memory consolidation. Nat Commun 14, 6100 (2023). 10.1038/s41467-023-41715-7

57 Rothe, T. et al. The Nuclear Receptor Nr4a1 Acts as a Microglia Rheostat and Serves as a Therapeutic Target in Autoimmune-Driven Central Nervous System Inflammation. J Immunol 198, 3878–3885 (2017). 10.4049/jimmunol.1600638

58 Sheng, R., Chen, C., Chen, H. & Yu, P. Repetitive transcranial magnetic stimulation for stroke rehabilitation: insights into the molecular and cellular mechanisms of neuroinflammation. Front Immunol 14, 1197422 (2023). 10.3389/fimmu.2023.1197422

59 Heck, C. N. et al. Two-year seizure reduction in adults with medically intractable partial onset epilepsy treated with responsive neurostimulation: final results of the RNS System Pivotal trial. Epilepsia 55, 432–441 (2014). 10.1111/epi.12534

60 Hansson, A. C. & Fuxe, K. Time-course of immediate early gene expression in hippocampal subregions of adrenalectomized rats after acute corticosterone challenge. Brain Res 1215, 1–10 (2008). 10.1016/j.brainres.2008.03.080

61 Cullinan, W. E., Herman, J. P., Battaglia, D. F., Akil, H. & Watson, S. J. Pattern and time course of immediate early gene expression in rat brain following acute stress. Neuroscience 64, 477–505 (1995). 10.1016/0306-4522(94)00355-9

62 Dobin, A. et al. STAR: ultrafast universal RNA-seq aligner. Bioinformatics 29, 15–21 (2013). 10.1093/bioinformatics/bts635

63 Liao, Y., Smyth, G. K. & Shi, W. featureCounts: an efficient general purpose program for assigning sequence reads to genomic features. Bioinformatics 30, 923–930 (2014). 10.1093/bioinformatics/btt656

64 Risso, D., Schwartz, K., Sherlock, G. & Dudoit, S. GC-content normalization for RNA-Seq data. BMC Bioinformatics 12, 480 (2011). 10.1186/1471-2105-12-480

65 Risso, D., Ngai, J., Speed, T. P. & Dudoit, S. Normalization of RNA-seq data using factor analysis of control genes or samples. Nat Biotechnol 32, 896–902 (2014). 10.1038/nbt.2931

66 Peixoto, L. et al. How data analysis affects power, reproducibility and biological insight of RNA-seq studies in complex datasets. Nucleic Acids Res 43, 7664–7674 (2015). 10.1093/nar/gkv736

67 Robinson, M. D., McCarthy, D. J. & Smyth, G. K. edgeR: a Bioconductor package for differential expression analysis of digital gene expression data. Bioinformatics 26, 139–140 (2010). 10.1093/bioinformatics/btp616

68 Butler, A., Hoffman, P., Smibert, P., Papalexi, E. & Satija, R. Integrating single-cell transcriptomic data across different conditions, technologies, and species. Nat Biotechnol 36, 411–420 (2018). 10.1038/nbt.4096

69 Hao, Y. et al. Integrated analysis of multimodal single-cell data. Cell 184, 3573–3587 e3529 (2021). 10.1016/j.cell.2021.04.048

70 Hao, Y. et al. Dictionary learning for integrative, multimodal and scalable single-cell analysis. Nat Biotechnol 42, 293–304 (2024). 10.1038/s41587-023-01767-y

71 Satija, R., Farrell, J. A., Gennert, D., Schier, A. F. & Regev, A. Spatial reconstruction of single-cell gene expression data. Nat Biotechnol 33, 495–502 (2015). 10.1038/nbt.3192

72 Stuart, T. et al. Comprehensive Integration of Single-Cell Data. Cell 177, 1888–1902 e1821 (2019). 10.1016/j.cell.2019.05.031

73. Rainer, J. (2017).

74 Zhang, Y. et al. Model-based analysis of ChIP-Seq (MACS). Genome Biol 9, R137 (2008). 10.1186/gb-2008-9-9-r137

75 Institute, A. Transcriptomic characterization of cell types in human M1 using droplet-based single nucleus RNA sequencing. (2022).

76 Eraslan, G., Simon, L. M., Mircea, M., Mueller, N. S. & Theis, F. J. Single-cell RNA-seq denoising using a deep count autoencoder. Nat Commun 10, 390 (2019). 10.1038/s41467-018-07931-2

77 Sciacca, M. L. a. K. G. a. E. glmmSeq: General Linear Mixed Models for Gene-Level Differential Expression. (2022).

78 International League Against Epilepsy Consortium on Complex, E. Genome-wide mega-analysis identifies 16 loci and highlights diverse biological mechanisms in the common epilepsies. Nat Commun 9, 5269 (2018). 10.1038/s41467-018-07524-z

79 Young, M. D., Wakefield, M. J., Smyth, G. K. & Oshlack, A. Gene ontology analysis for RNA-seq: accounting for selection bias. Genome Biol 11, R14 (2010). 10.1186/gb-2010-11-2-r14

80 Paczkowska, M. et al. Integrative pathway enrichment analysis of multivariate omics data. Nat Commun 11, 735 (2020). 10.1038/s41467-019-13983-9

81 Milacic, M. et al. The Reactome Pathway Knowledgebase 2024. Nucleic Acids Res 52, D672–D678 (2024). 10.1093/nar/gkad1025

