## Supplemental information for "The gene expression signature of electrical stimulation in the human brain"

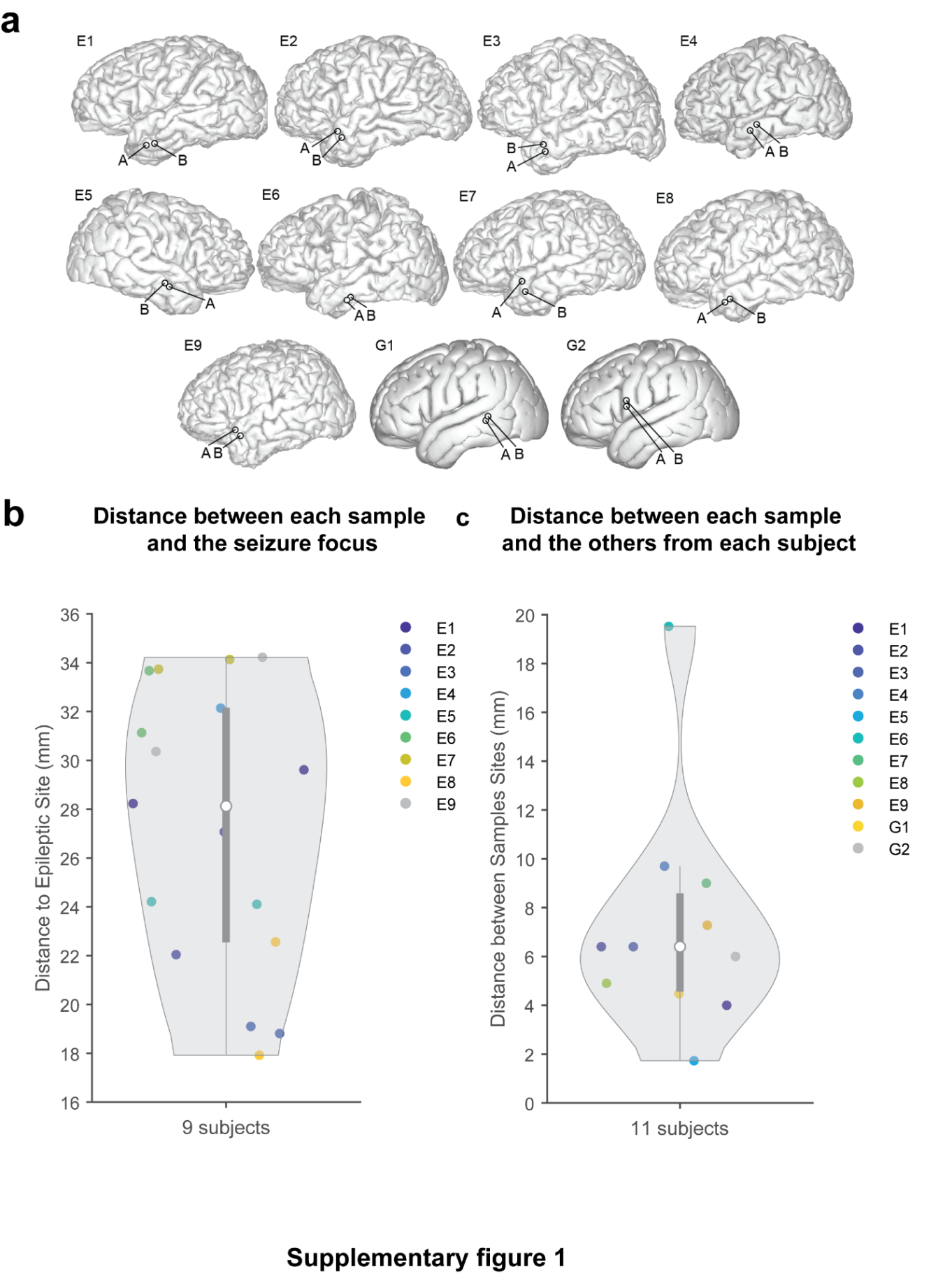

**Supplementary Figure 1. Participant-specific resection sample location. a.** Reconstructed brain images using preoperative T1 sequences. Intraoperative photos of the sample sites were obtained and mapped onto brain reconstructions based on anatomical landmarks. Site A: baseline, Site B: stimulated or unstimulated samples (30 mins after baseline). **b.** The distance (in mm) between each resection sample and the border of the seizure focus in the epilepsy group. **c.** Distance (in mm) between the two resection samples (A and B), plotted for all sub-sets.

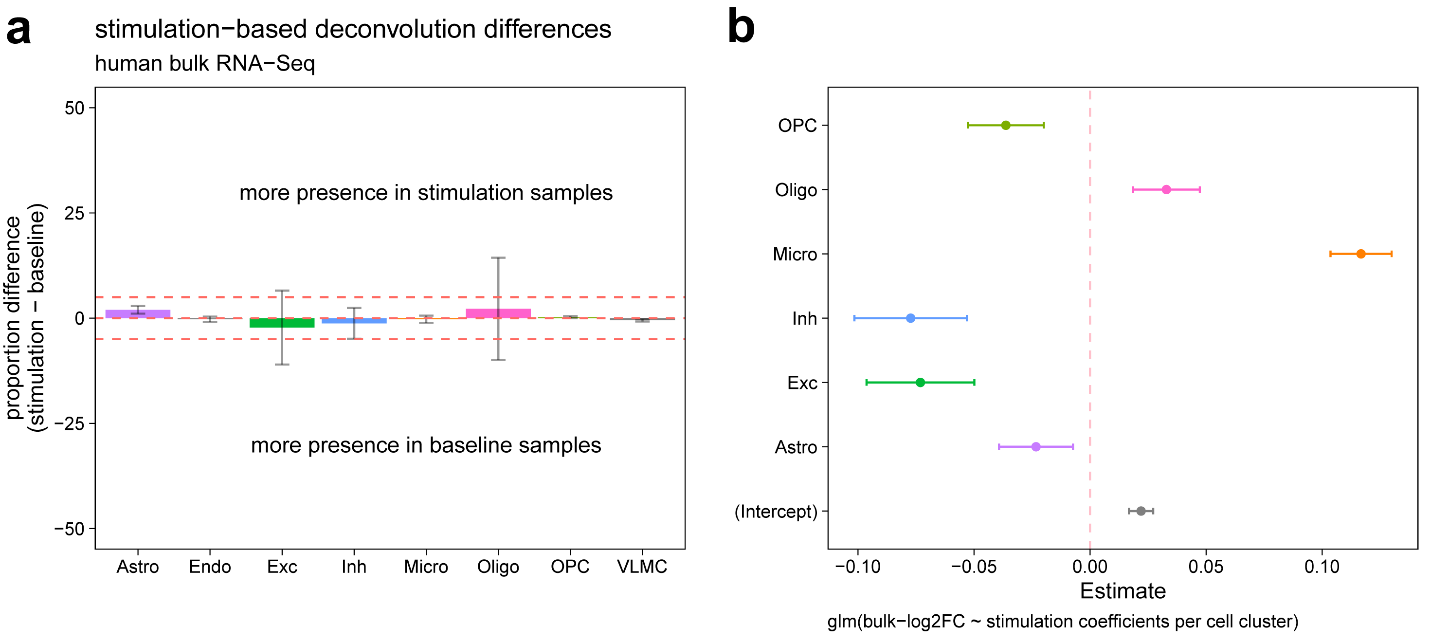

**Supplementary Figure 2. Cell-type deconvolution of the human bulk RNA-seq using single nuclei RNA-seq data.** Bar plots showing the mean change in cell type proportions between stimulated and baseline samples from the stimulated paradigm. Each point represents a participant-level difference in estimated proportion (stimulated – baseline) for a given cell type. Horizontal dashed red lines represents a threshold of ±5%.

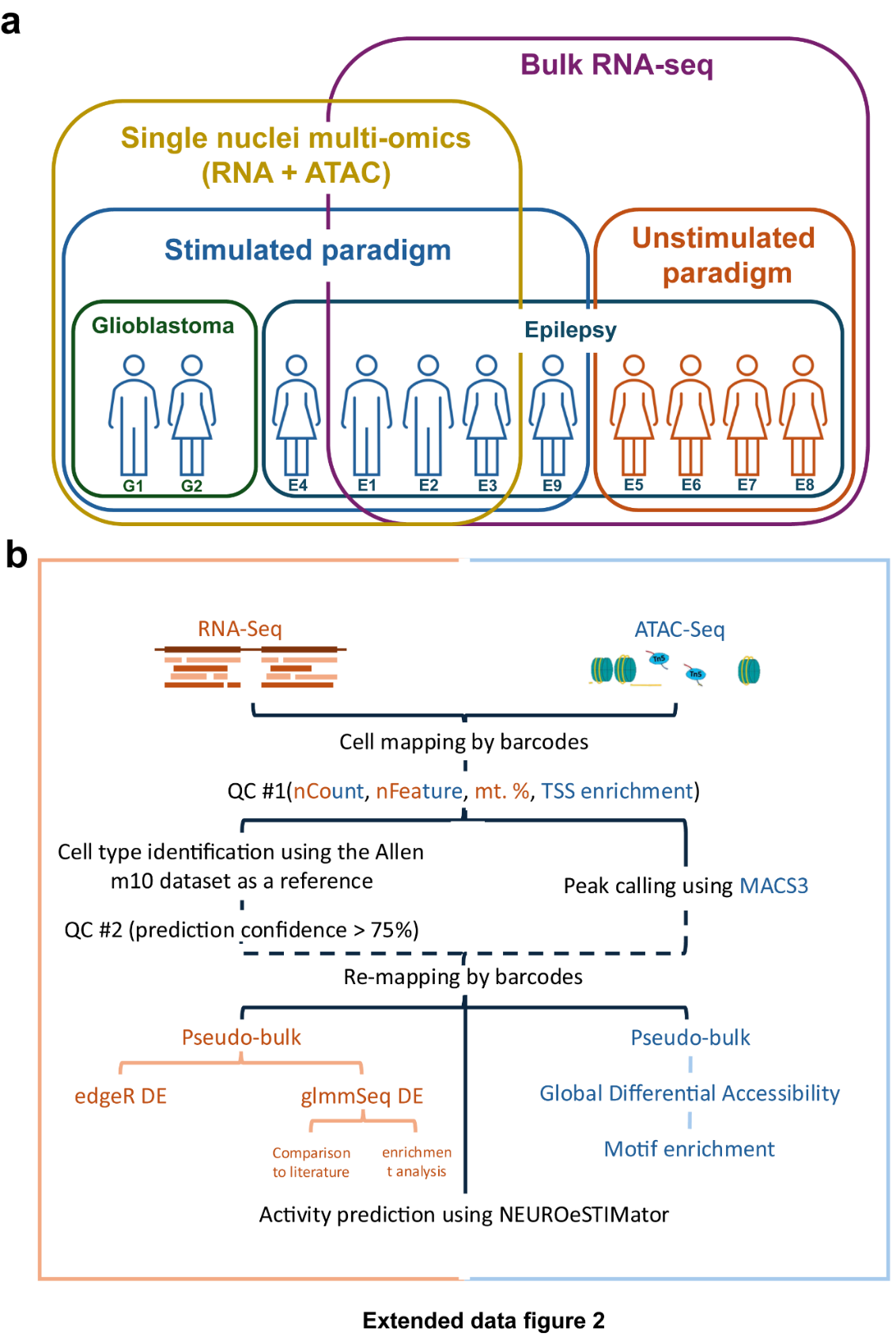

**Supplementary Figure 3. a.** Samples for bulk RNA and single nuclei multiomics. **b.** Analysis pipeline for the single nuclei multi-omics data.
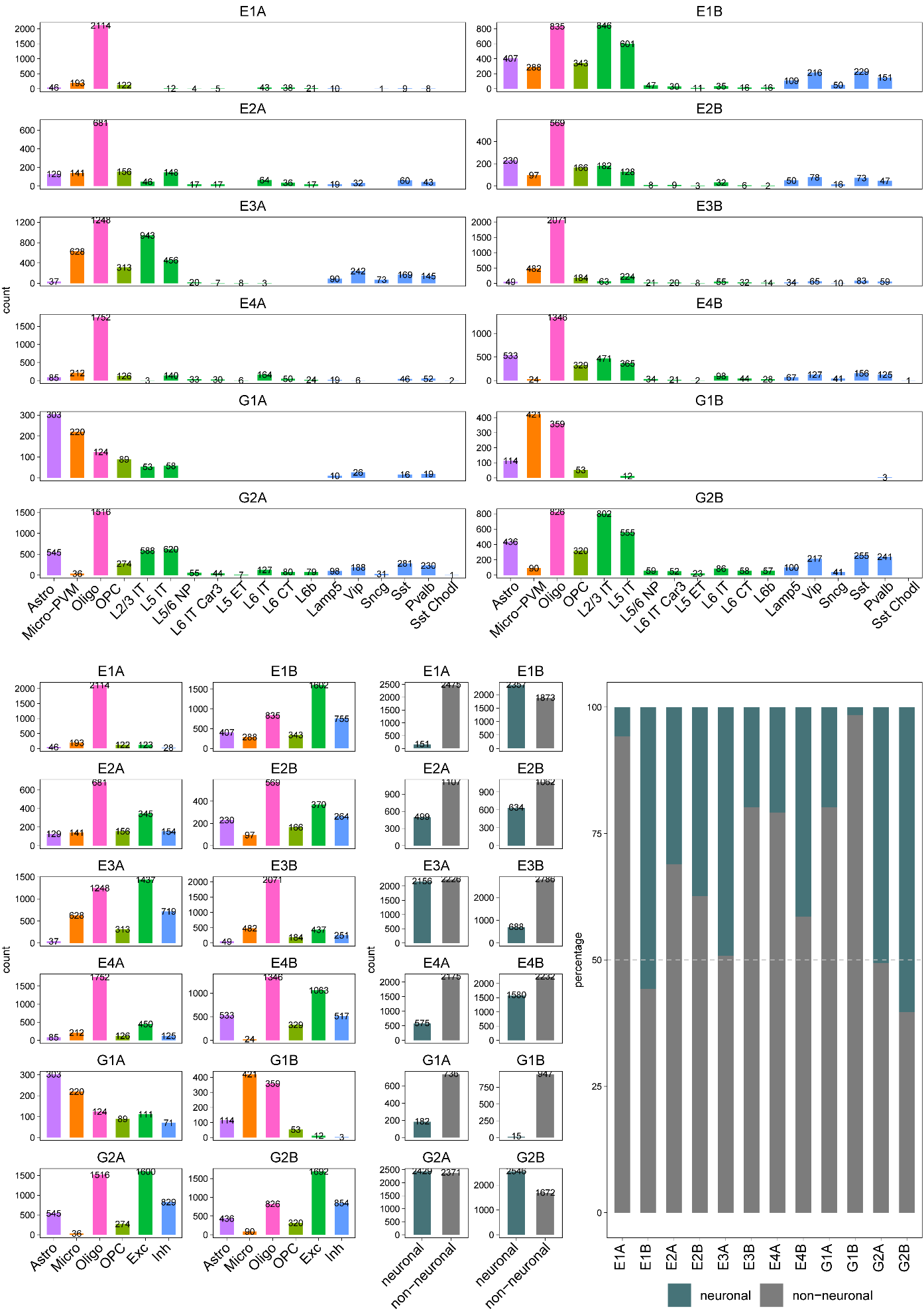
**Supplementary Figure 4. Composition of cell types in single-nuclei multi-omics samples.** **a,** Bar plots showing the distribution of annotated cell types (including neuronal subtypes) across six participants, each with baseline and corresponding stimulated samples. **b,** Counts of major cell type clusters aggregated into excitatory neurons, inhibitory neurons, microglia, astrocytes, oligodendrocytes, and oligodendrocyte precursor cells (OPCs). **c,** Total counts of neuronal (excitatory and inhibitory) versus non-neuronal (microglia, astrocytes, oligodendrocytes, OPCs) cells per sample. **d,** Ratio of neuronal to non-neuronal cells for each sample, highlighting variation in cellular composition across patients and conditions.

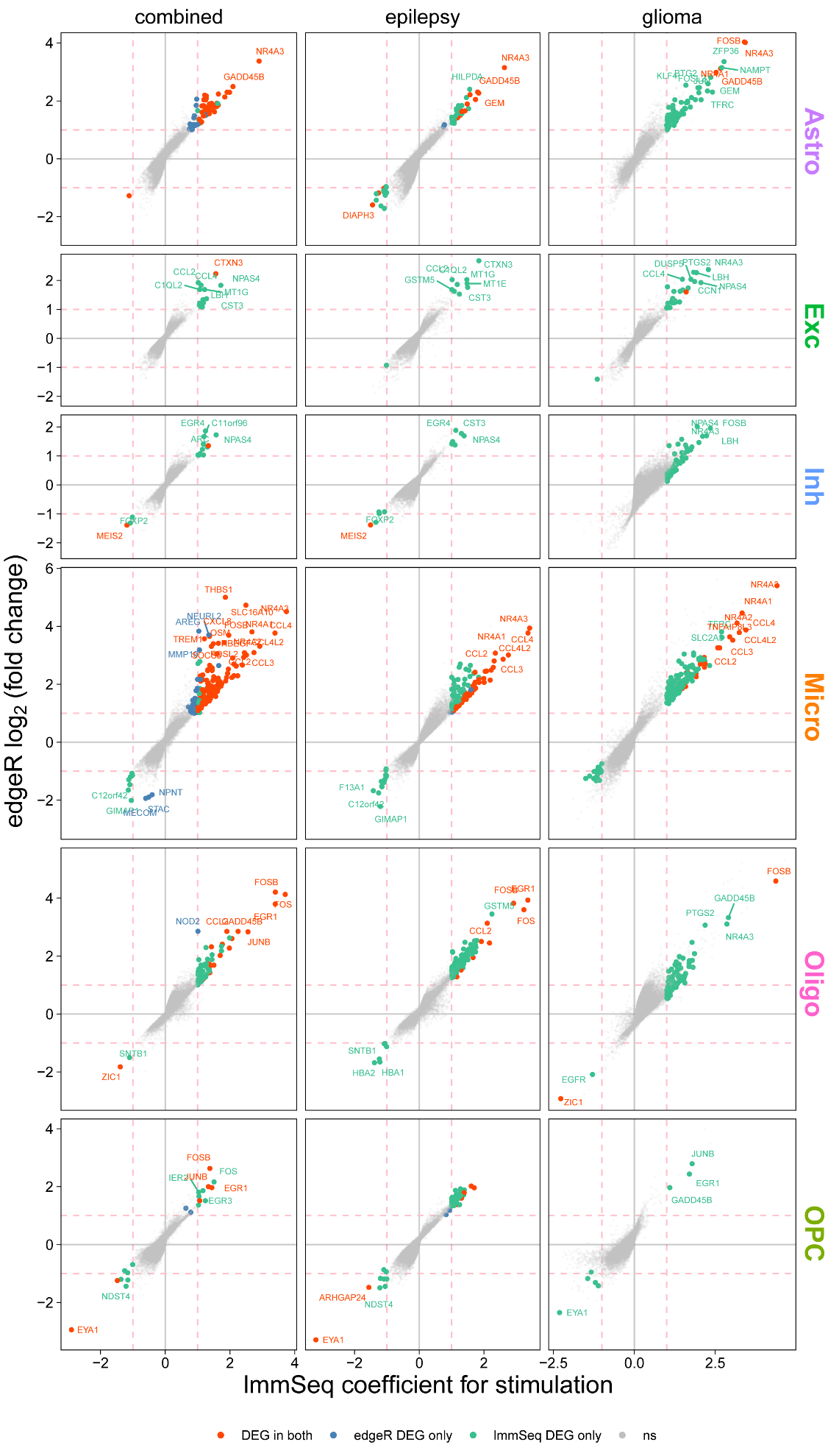

**Supplementary Figure 5.** **Comparison of eSTIM-induced differential gene expression signature.** Scatter plots comparing differential gene expression estimates from two modeling approaches (edgeR and lmmSeq) across six major brain cell types. The x-axis represents the stimulation coefficient estimated using a linear mixed-effects model, which accounts for participant-level random effects, while the y-axis shows the log₂ fold change of the same genes estimated using the edgeR model. Analyses were performed using three group sets: epilepsy samples only (n=4 pairs of samples), glioblastoma samples only (n=2 pairs of samples), and all samples combined (n=6 pairs of samples). The edgeR model for the third group (all samples combined) included diagnosis (epilepsy/glioblastoma), age, sex, and anesthesia as covariates. The mixed-effects model for the same group included the same covariates with the addition of the participant ID as a random variable. Colored points highlight differentially expressed genes, where red points represent genes identified as DEGs in both models (lmmSeq and edgeR), blue points represent DEGs identified by edgeR and not lmmSeq, green points represent DEGs identified by lmmSeq and not edgeR, and grey points show genes that were not differentially expressed as indicated by both models. Genes with |log2(FC)| > 1 and FDR < 0.05 were considered differentially expressed in edgeR results. Genes with |stimulation beta coefficient| > 1 and FDR < 0.05 in lmmSeq results were considered differentially expressed.

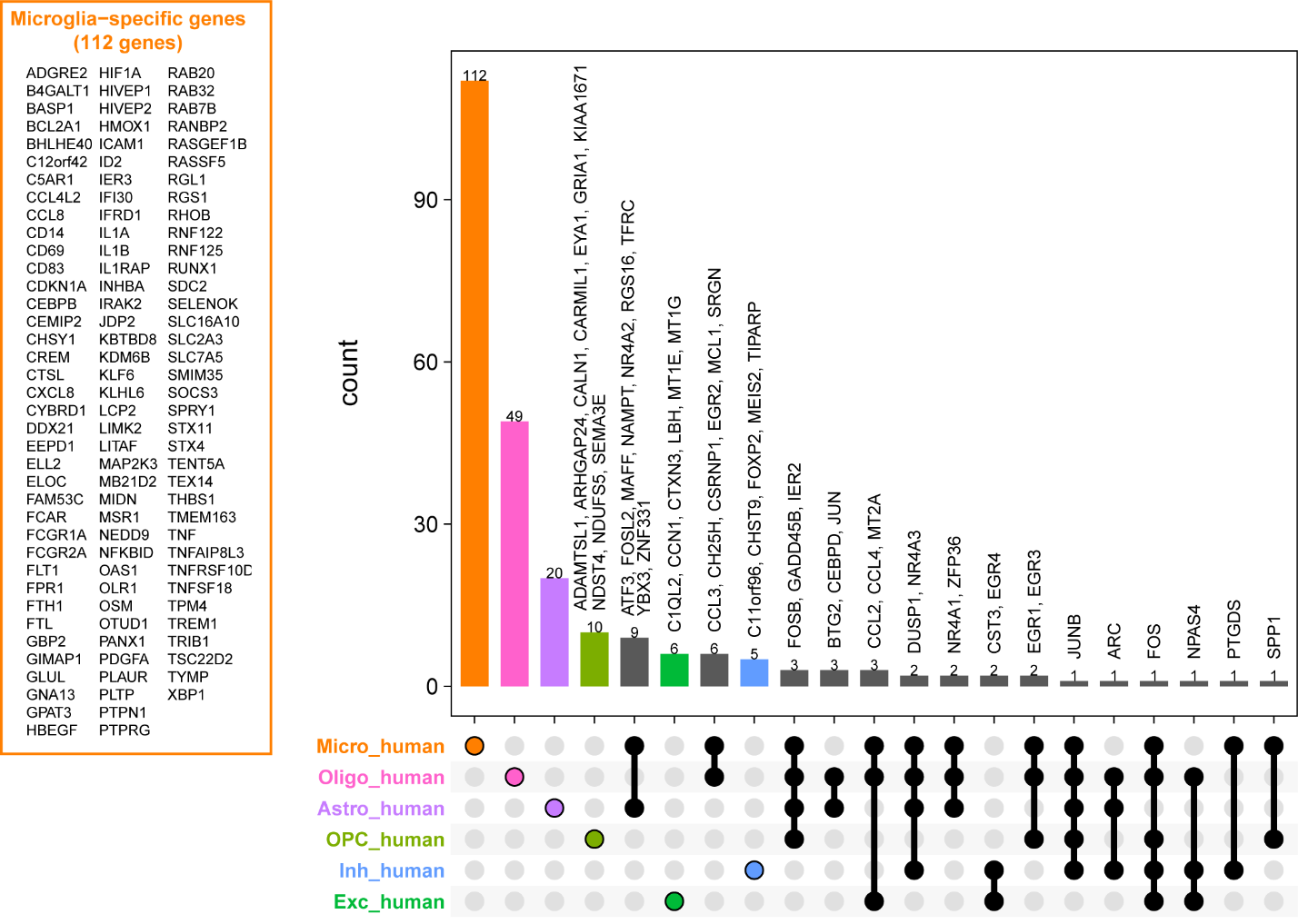

**Supplementary Figure 6.** **Upset plot showing the unique and overlapping genes induced by eSTIM across different cell types from snRNA-seq data.**

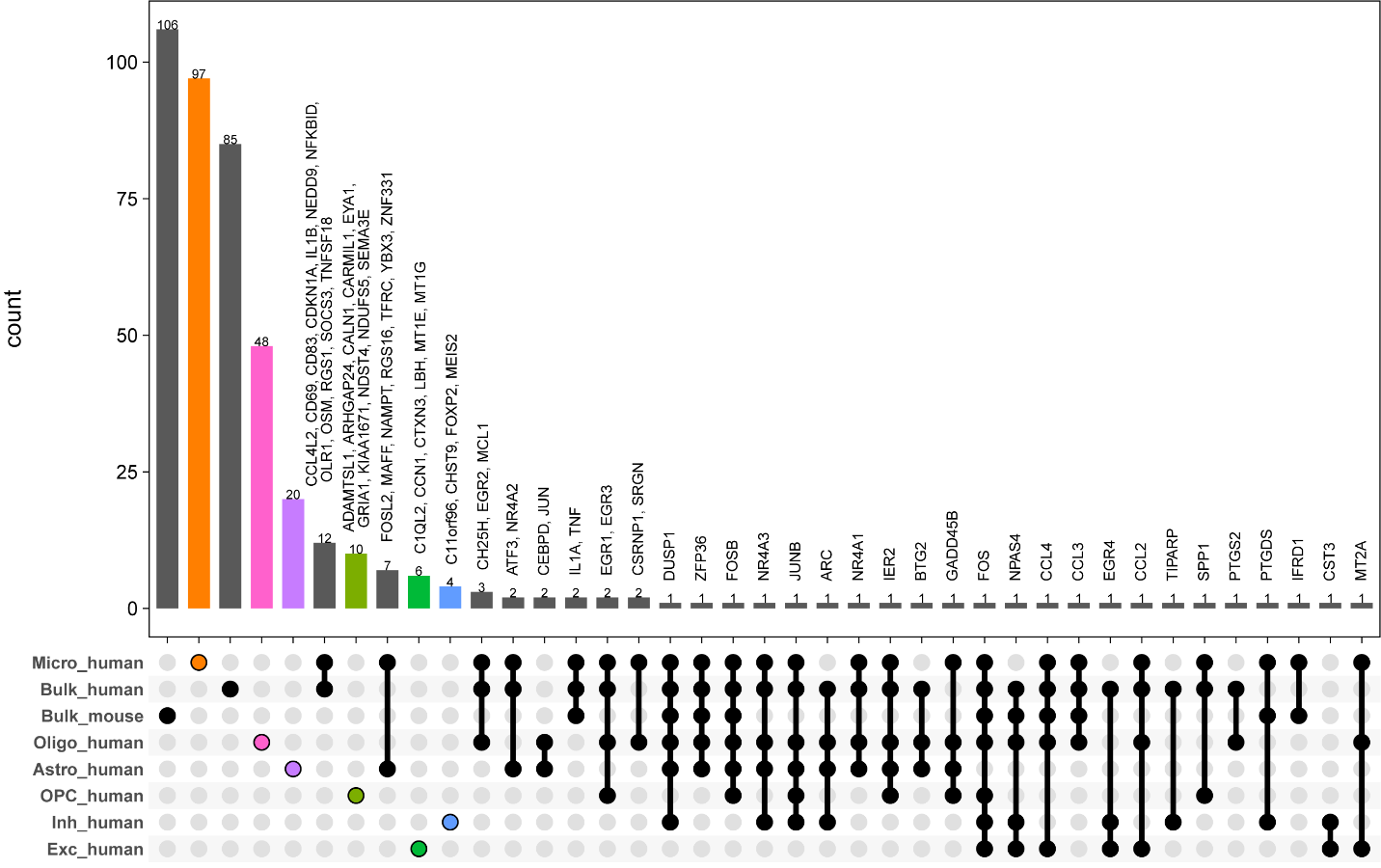

**Supplementary Figure 7.** **Upset plot comparing DEGs from snRNA seq and bulk RNA seq following eSTIM.** DEGs from human bulk RNA-Seq data from the stimulation paradigm and snRNAseq from the stimulation paradigm are illustrated in an Upset plot. For human bulk RNAseq and mouse bulk RNA seq, genes with |log2(FC)| > 0.2 and FDR < 0.05 were considered differentially expressed. For single nuclei RNA seq, |stimulation beta coefficient| > 1 and FDR < 0.05 were considered differentially expressed for this analysis.

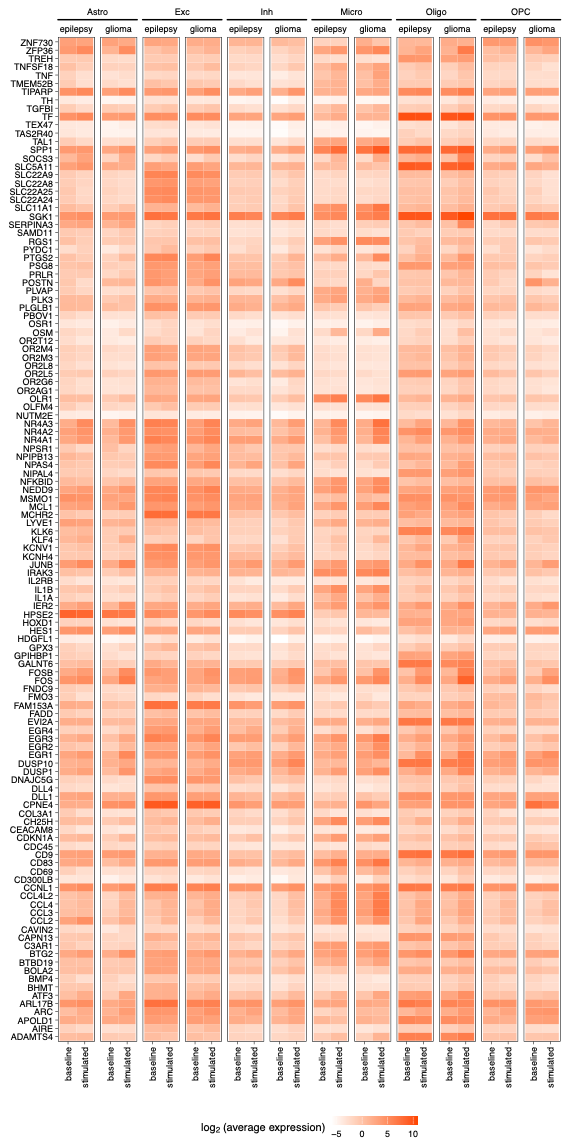

**Supplementary Figure 8.** **Expression profiles of eSTIM-induced genes from bulk RNA seq and snRNAseq.** A heatmap showing the expression of significant DEGs identified in human bulk RNA-seq within human snRNA-seq samples. Expression levels are shown separately for stimulated and unstimulated conditions across major brain cell types, divided by diagnosis group (epilepsy and glioblastoma).

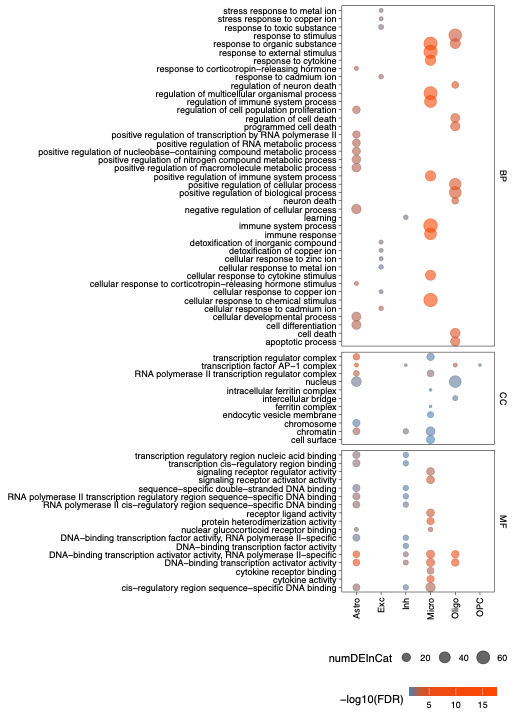

**Supplementary Figure 9.** **Gene ontology (GO) enrichment of eSTIM-induced DEGs across major cell types from snRNAseq data.** Dot plot showing enriched Gene Ontology (GO) terms among DEGs across major brain cell types. Each point represents a significant GO term (FDR < 0.05), with the circle size indicating the number of DEGs annotated to the term. BP: Biological Processes. CC: Cellular Component. MF: Molecular Function.

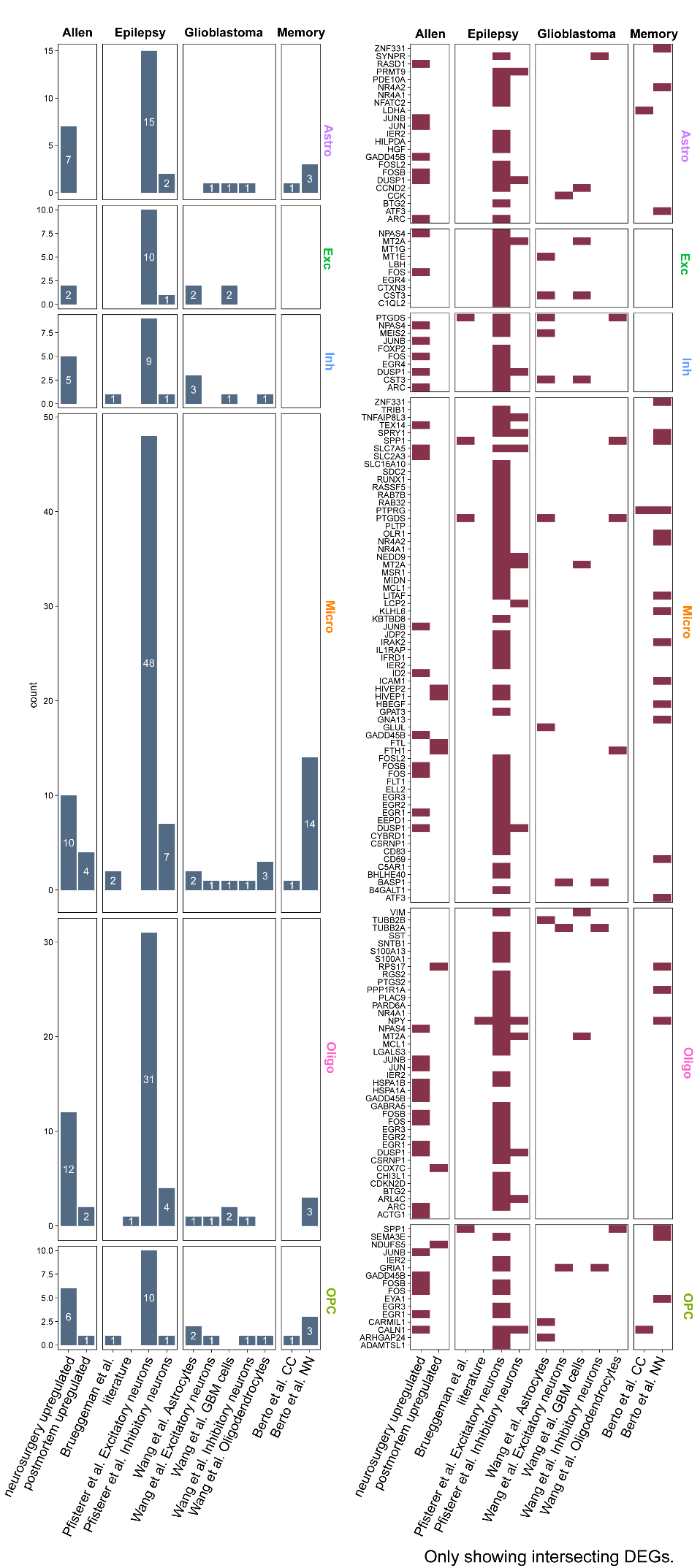

**Supplementary Figure 10.** **Overlap of eSTIM-responsive genes with disease- and trait-associated genes from the literature.** Bar plot showing the overlap between DEGs identified in each major brain cell type and curated external datasets, including epilepsy-associated genes (from GWAS, prior literature, and transcriptomics), glioblastoma single-cell RNA-seq data, memory-related gene sets, and neurosurgery/postmortem transcriptomic datasets.

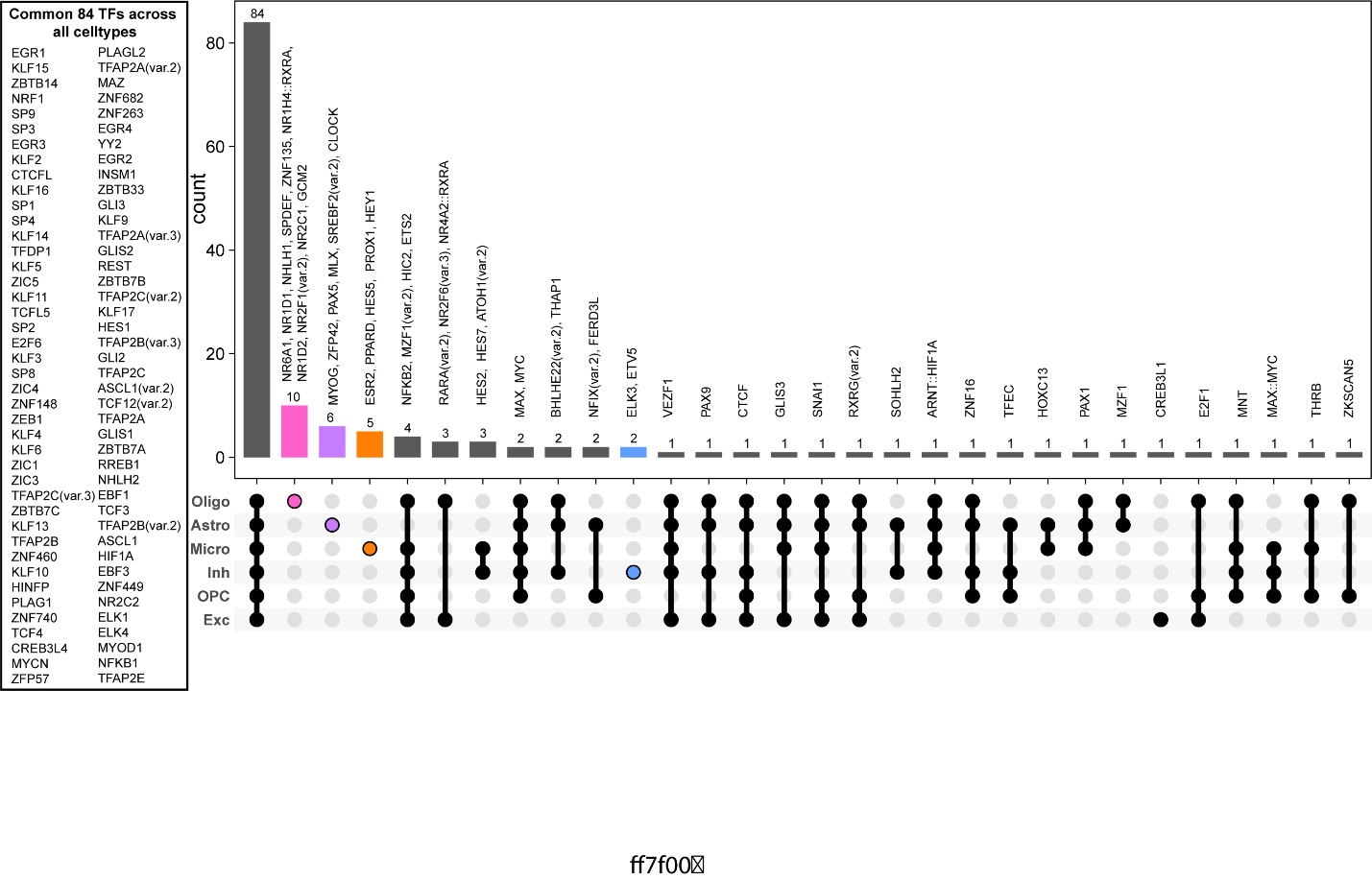

**Supplementary Figure 11.** **Upset plot showing the distinct and overlapping TF motifs enriched following eSTIM in the snATAC-seq.**

**Supplementary Table 1 | Participant metadata.** Clinical and demographic information for all 11 neurosurgical patients included in the study. The table includes participant ID, sex, age, clinical condition (medically refractory epilepsy or glioblastoma), anesthesia status (awake or sedated), and availability of neuropsychological assessment. Samples are also annotated by their experimental paradigm (stimulated or unstimulated).

**Supplementary Table 2 | Bulk RNA-seq analysis of stimulated samples (Human).** Differentially expressed genes (DEGs) from bulk RNA-seq comparing electrically stimulated (eSTIM) samples to their matched baseline.

**Supplementary Table 3 | Bulk RNA-seq analysis of unstimulated control samples (Human).** Differentially expressed genes (DEGs) from bulk RNA-seq comparing the second, unstimulated sample to the initial baseline sample in the unstimulated paradigm.

**Supplementary Table 4 | Bulk RNA-seq analysis of down-stream effects (Mouse).** Differentially expressed genes (DEGs) from bulk RNA-seq comparing post-stimulation samples from the stimulated paradigm to baseline samples from the unstimulated paradigm, identifying downstream transcriptional effects.

**Supplementary Table 5 | Multi-omics single-nuclei RNA-seq differential expression results (Human).** Differential gene expression analysis performed on a pseudobulk level for each major cell type. The Excel file contains separate sheets for excitatory neurons, inhibitory neurons, microglia, and other identified cell types. The first sheet provides a detailed explanation of each column header used across all sheets.

**Supplementary Table 6 | Gene Ontology (GO) enrichment for differentially expressed genes (Human).** Significantly enriched Biological Processes, Molecular Functions, and Cellular Components from Gene Ontology analysis performed on the differentially expressed genes identified per cell type from the multi-omics snRNA-seq data.

**Supplementary Table 7 | Pathway enrichment analysis for differentially expressed genes (Human).** Results from pathway enrichment analysis (Reactome) performed on the differentially expressed genes identified per cell type, highlighting significant enrichment in cytokine and chemokine-related pathways.

**Supplementary Table 8 | Transcription factor motif enrichment analysis (Human).** Results from the transcription factor binding motif enrichment analysis performed on accessible chromatin regions from multiple cell types. The table details significantly enriched motifs, including those for the EGR sub-family, across different cell populations.
